# Genetic control of leaf carbon- and water-related traits in grapevine under greenhouse and field conditions

**DOI:** 10.64898/2026.10.02.754987

**Authors:** Eva Coindre, Sarah Désirée Etou, Romain Boulord, Agnès Doligez, Thierry Simonneau, Nicolas Saurin, Gautier Sarah, Baptiste Pierre, Roberto Bacilieri, Dominique This, Benoît Pallas, Vincent Segura, Aude Coupel-Ledru

## Abstract

Understanding the genetic architecture of ecophysiological traits related to carbon gain and water loss is crucial to develop cultivars tolerant to abiotic stresses and that maintain productivity in the context of climate change. This can be achieved both through experiments in the greenhouse on potted plants under controlled conditions and in the field. However, the stability of genetic determinism of ecophysiological traits across environmental conditions has rarely been evaluated. We studied a grapevine diversity panel grown both in a greenhouse and in the field, to investigate the impact of growing conditions on leaf ecophysiological traits related to leaf morphology, fluorescence, gas exchange and water use efficiency. The correlation structure between traits was conserved within each environment even though, for most traits, cultivars ranking was not conserved between both environments. However, a set of cultivars, sharing the same functioning pattern in terms of gas exchange and fluorescence in both environments, was identified. Single-and multi-environment Genome-Wide Association Studies (GWAS), based on a dense genotyping matrix, were used to dissect the genetic architecture of leaf traits. The Quantitative Trait Loci (QTLs) identified through single-environment analyses differed substantially from those detected using multi-environment ones. Multi-environment approaches highlighted shared QTLs across environments for gas exchange and fluorescence traits. Candidate genes associated with carbon- and water-related functions were further identified in these genomic regions. Despite the weak correlation between greenhouse and field trait values, combining both environments enabled the identification of robust genetic regions associated with ecophysiological traits of interest.

## Introduction

Climate change is a major challenge impacting crop production. One major criterion determining plant adaptation to stressful conditions is the balance between carbon gain through net CO_2_ assimilation (A_n_), and water loss through transpiration (E) or stomatal conductance (g_s_). Estimating water use efficiency (WUE) as the ratio between A_n_ and E (instantaneous WUE) or g_s_ (intrinsic WUE), is relevant because it integrates these processes and provides an estimate of the plant’s ability to balance water loss into carbon gain. Selecting cultivars with high WUE, without compromising growth and production, is a key target for adapting crops to climate change. At the leaf level, A_n_, E and g_s_ can be measured directly using gas exchange analyzers which allow a precise and non-destructive measurement of WUE (Coupel-Ledru, 2021). WUE is also influenced by leaf morphological traits such as leaf mass per area (LMA) or leaf water content (WC_f_), which can modify water status, CO_2_ diffusion and photosynthetic activity.

However, characterizing the genetic determinism of these traits is difficult, particularly because high-throughput phenotyping of hundreds or thousands of plants within a short time frame remains challenging. g_s_ and E can be easily and quickly measured at the leaf level using new generation porometer devices. However estimating A_n_ requires a gas exchange analyzer which substantially increases the time required for phenotyping. To overcome this limitation, chlorophyll fluorescence parameters, related to the efficiency of Photosystem II (ΦPSII) and electron transport rate in the photosynthetic chain (ETR), have been suggested as key traits to provide information on leaf photosynthetic activity. They were used in multivariate models, together with leaf temperature in fruit trees (Losciale et al., 2015) or with g_s_, E and microclimatic variables measured with a new generation porometer in grapevine (Coindre et al., 2026) to predict A_n_. Measuring leaf mass per area (LMA) or leaf water content (WC_f_) is also time-consuming, as they require manual weighing of leaves. As an alternative, Near InfraRed Spectroscopy (NIRS) has been reported to provide valuable predictions of LMA and WC_f_ (Ge et al., 2019; Coindre et al., 2026). These recent advances in phenotyping methods facilitate the study of the genetic determinism of carbon- and water-related leaf traits in large populations.

In this context, high-throughput phenotyping platforms are particularly useful for assessing traits of interest for hundreds of genotypes under controlled and consistent atmospheric and soil conditions for each plant, thereby minimising environmental bias when comparing genotypes. In particular, they enable reproducible monitoring of soil water deficit intensities, allowing a fine characterisation of the genotype response to soil water deficit (Cabrera-Bosquet et al., 2016; Coupel-Ledru et al., 2014). However, the environmental conditions applied in platform experiments generally correspond to a more limited range of variation than those observed outdoors, particularly in terms of temporal and spatial variability of environmental and soil conditions. Thereby, extrapoling results from one environment to another is challenging, especially from greenhouse to field under agronomic conditions (Poorter et al., 2016). Under field conditions, multiple biotic and abiotic constraints interact, which can lead to more unexplained variability than in controlled conditions. Leaf traits are affected by environmental conditions, such as the amount of intercepted solar radiation (Wimalasekera, 2019) and field environments are generally characterized by higher evaporative demand and light intensity than greenhouse conditions. Moreover, the use of potted plants as opposed to field-rooted plants, is known to limit plant growth by decreasing photosynthesis and affecting root development (Poorter et al., 2012). Contrasted results have been reported in studies comparing multiple environments for leaf carbon and water related traits. For instance in a germplasm of 76 genotypes of wheat, Silva-Pérez et al., 2020 found high genetic correlations between greenhouse and field environments for A_n_, chlorophyll index (measured with SPAD) and leaf nitrogen concentration, while for the same species with 80 wheat lines, Sales et al., 2022 did not find any correlation at all between environments for photosynthetic traits. In switchgrass, large differences in coefficients of variation for A_n_ and g_s_ were reported between greenhouse and shelter environments for the same genotype (Lovell et al., 2016). Conversely, high consistency has been reported for yield components, measured in greenhouse and field conditions. For instance a study on 19 maize hybrids reported that the genotypes with the most sensitive leaf elongation rate to water deficit in the greenhouse were the ones that most reduced seed number under water deficit in the field (Chapuis et al., 2012). In a larger panel of maize hybrids, the correlation between leaf number and thermal time since emergence was similar whether the experiment was performed in a field or a greenhouse environment (Millet et al., 2019). Nevertheless, these traits may be more conservative and less sensitive to environmental variation than leaf functioning traits. The problem of translating results from greenhouse to field is even more difficult with woody perennials that are generally much smaller and younger in the greenhouse than in the field. Consequently, to the best of our knowledge, no quantitative genetic study has been carried out on both greenhouse and field environments to assess the stability of detected Quantitative Trait Loci (QTLs) in perennial crops. This study focuses on cultivated grapevine, *Vitis vinifera* L. subsp*. vinifera* (hereafter referred to as *Vitis vinifera*), which is one of the most economically important horticultural crops (Alston & Sambucci, 2019). Breeding programs for this perennial crop can last for more than 20 years (Töpfer & Trapp, 2022). Therefore, there is an urgent need to reduce the selection duration by enabling marker assisted selection or genomic prediction (Avia et al., 2023). Nevertheless, the integration of these methods into breeding programs requires deciphering the genetic determinism of adaptative traits. In grapevine, a few studies already found QTLs for water related traits in greenhouse environments (Coupel-Ledru et al., 2014, 2016; Marguerit et al., 2012), and initiated comparison with vineyard conditions (Coupel-Ledru et al., 2022) but they focused on biparental populations and thus on a fairly restricted genetic diversity. We recently reported a QTL related to g_s_ in the field in a large diversity panel (Chir et al., 2025). However, the consistency of the QTLs effects across greenhouse and field environments for multiple leaf traits has never been investigated.

In this study, we compared carbon- and water-related traits obtained on a grapevine diversity panel (Nicolas et al., 2016), grown in pots in a greenhouse and under field conditions. We performed single- and multi-environment Genome-Wide Association Studies (GWAS) to characterize the genetic determinism of these traits and specifically look for stable QTLs across environments.

## Results

### Variation of genotypic values of ecophysiological traits in greenhouse and field environments

Ecophysiological traits were characterized for a diversity panel of 279 cultivars in both a greenhouse and a field environment. Some of the traits (Table S1) were directly measured, in both environments, including SPAD, electron transport rate (ETR), quantum efficiency of photosystem II in light (ΦPSII), stomatal conductance (g_s_) and transpiration rate (E). Leaf mass per area (LMA), water content (WC_f_) and water quantity (WQ) were directly measured in the greenhouse while Partial Least Square Regression (PLSR) models based on Near InfraRed Spectroscopy (NIRS) were used to predict them in the field (Table S2, R²_val_ > 0.80 for LMA and WC_f_, and R²_val_ around 0.6 for WQ, Table S3). Net CO_2_ assimilation rate (A_n_) was predicted in both environments using a PLSR model based on poro-fluorometry measurements (R²_val_ = 0.67 in the field, Table S3 and R²_val_ = 0.80 in the greenhouse (Coindre et al., 2026)).

In each environment, we used linear mixed-models to estimate genotypic values (BLUPs) for all the traits. The selected models always retained the fixed effect of the date of measurement. Fixed effects of climatic variables, such as air temperature (T_air_), leaf vapor pressure deficit (VPD) and photosynthetically photon flux density (PPFD), were retained for g_s_, E, ΦPSII, ETR, A_n_ and WUE_intr_cal_ for the field environment, but not for the greenhouse environment (Table S4). This may result from a higher variability in climatic conditions in the field compared to greenhouse (Figure S1).

We observed a large variability in BLUPs for each trait in each environment, as well as clear differences in mean values between the greenhouse and the field (Figure 1). SPAD, WC_f_ and ΦPSII displayed higher values in the greenhouse while LMA, WQ, ETR, g_s_, E, A_n_, and WUE_intr_ca_ were higher in the field (Figure 1). Among all traits, A_n_ showed the largest difference between environments, increasing by 2.5-fold from greenhouse (mean 6.2 µmol m⁻² s⁻¹) to field (mean 15.4 µmol m⁻² s⁻¹) while g_s_ increased only by 1.5-fold. This resulted in a higher WUE_intr_cal_ in the field than in the greenhouse. ETR was more affected by the environment (1.7-fold increase) than ΦPSII (1.3-fold decrease). This greater difference in ETR likely reflects differences in PPFD between environments, as PPFD is directly incorporated into ETR calculation (Figure S1a). The range of variation within BLUPs values was similar between environments for most traits, except for ΦPSII, g_s_, E, A_n_ and WUE_intr_cal_ where there was less genotypic variability in the field compared to the greenhouse (Figure 1f, g, h and i).

**Figure 1:**
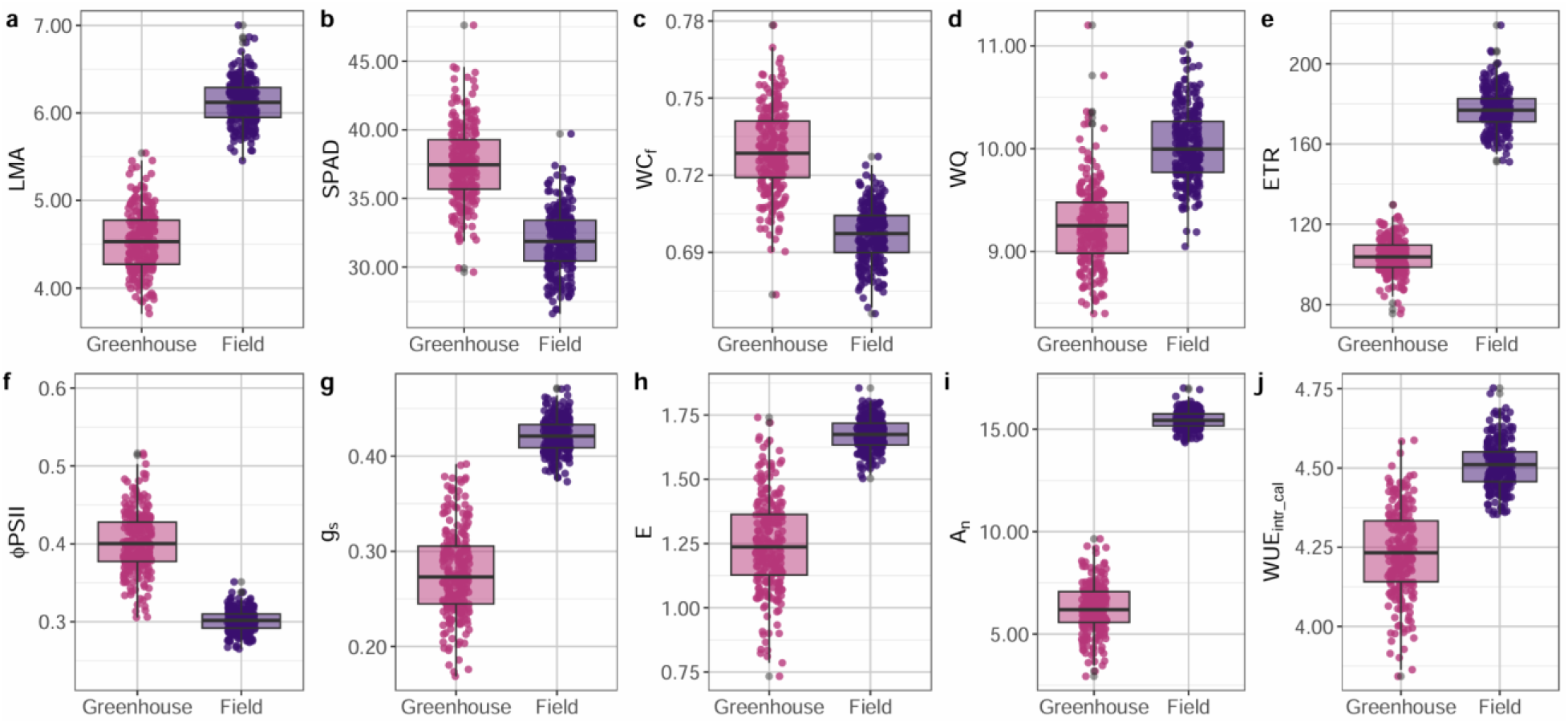
Boxplots of genotypic values (BLUPs) for carbon- and water-related leaf traits in greenhouse and field environments. **a** Leaf mass per area (LMA in mg cm^-^²), **b** Proxy of chlorophyll content (SPAD), **c** Water content (WCf in mg H2O mg^-1^ fresh weight), **d** Water quantity (WQ in mg H2O per leaf disk), **e** Electron transport rate (ETR in µmol m^-2^ s^-1^), **f** Quantum efficiency of photosystem II in light (ΦPSII), **g** Stomatal conductance (gs in mol m^-2^ s^-1^), **h** Transpiration rate (E in mmol m^-2^ s^-1^), **i** Net CO2 assimilation rate (A_n_ in µmol m^-2^ s^-1^) and **j** Intrinsic water use efficiency (WUE_intr_cal_ in µmol CO_2_ mol^-1^ H_2_O). Square-root transformation was applied to WQ, g_s_ and E and log transformation (base 10) was applied to WUE_intr_cal_ before adjusting the linear mixed-model, as the data were not normally distributed. n = 246 cultivars in the greenhouse environment and n = 255 cultivars in the field environment.

### Intra- and inter-environment genetic correlations between traits

The structure of correlations between genotypic traits values was broadly similar within both environments (Figure 2a). g_s_ and E were strongly correlated (*r* > 0.98) together, and significantly correlated with ETR and ΦPSII (r > 0.39). These four traits were also significantly and positively correlated with A_n_ (Figures 2a and S2). WUE_intr_cal_ was negatively correlated with A_n_ and g_s_ (r < - 0.45), with a stronger correlation with g_s_, meaning that WUE_intr_cal_ was more influenced by water losses than by CO_2_ assimilation, whatever the environment. ETR and ΦPSII were the only traits showing distinctly different correlations between environments, with a much stronger correlation in the field (r = 0.91) than in the greenhouse (r = 0.37) (Figure 2a, Figure S2). Regarding the other traits, there was a significant, medium correlation between SPAD and A_n_ with an r of 0.29 and 0.37 in greenhouse and field environments, respectively. In both environments, SPAD was also positively correlated with LMA (r = 0.54) and LMA with WQ (r > 0.61). The traits that most strongly structured the observed variability were A_n_, E, g_s_, WUE_intr_cal_, ETR and ΦPSII, which contributed to the first component of the Principal Component Analyses (PCA). This first component explained more than 42.7% of the genotypic variability, regardless of the environment (Figure 2b and c). LMA, WQ, SPAD and WC_f_ contributed less to the observed variability, as they were mainly involved in the second PCA component, which explained no more than 20.1% of genotypic variability.

**Figure 2:**
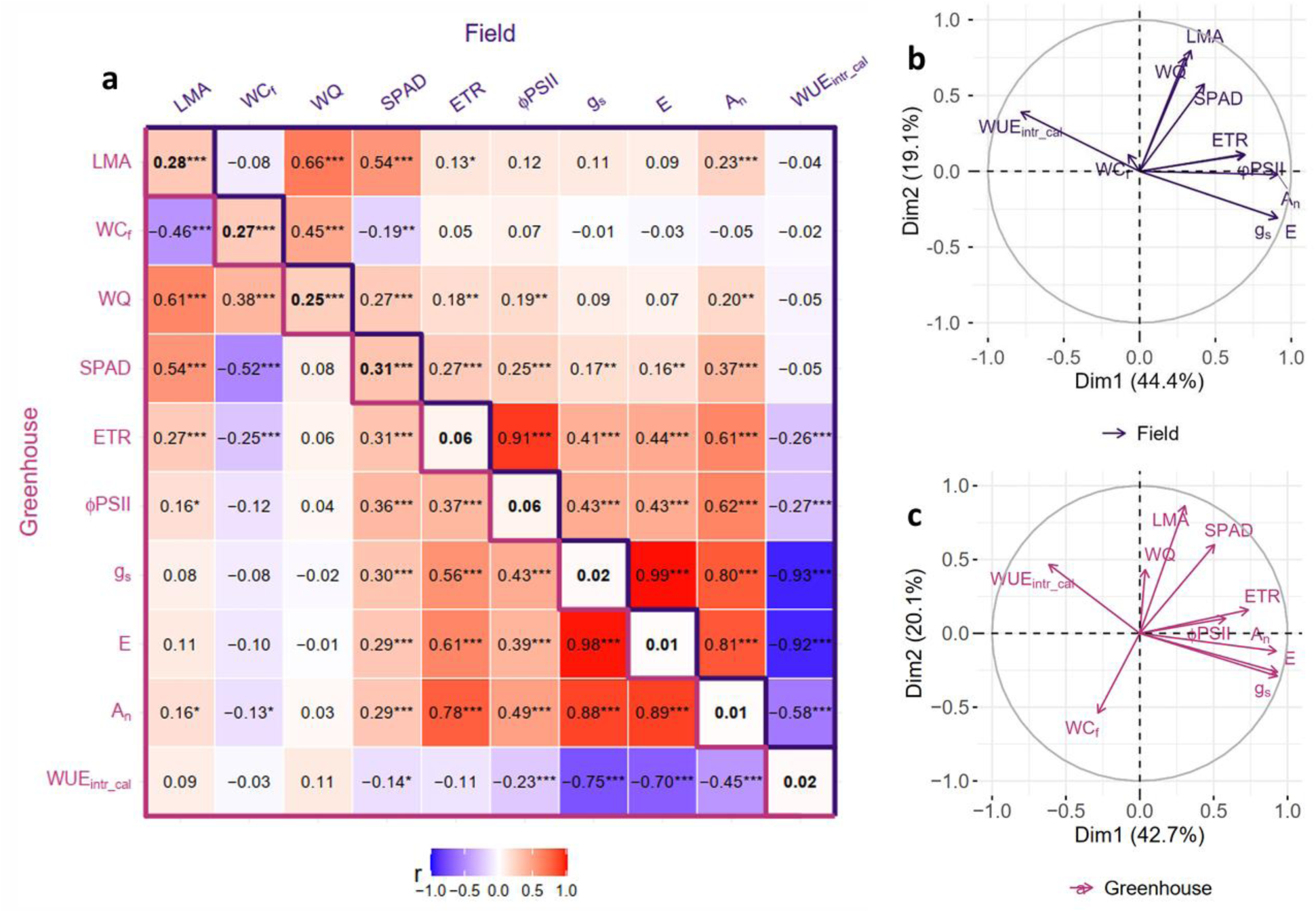
Genetic correlations between traits across environments and Principal Component Analyses (PCA) per environment. **a** Matrix of Pearson correlation coefficients between BLUPs of all carbon- and water-related traits in each environment (upper part highlighted in purple: field environment with n = 255 cultivars, lower part highlighted in pink: greenhouse environment with n = 246 cultivars). The diagonal indicates the correlation coefficients between the two environments for each individual trait (n = 222 cultivars). Significance level is indicated for each correlation as follows: * = *P*-value ≤ 0.05; ** = *P*-value ≤ 0.01; *** = *P*-value ≤ 0.001. **b** and **c** Plots of trait contribution to the two principal components of the PCA, for the field (n = 255 cultivars) (b) and greenhouse (n = 246 cultivars) (c) environments, respectively. LMA: Leaf mass per area, SPAD: Proxy of chlorophyll content, WC_f_: Water content, WQ: Water quantity, ETR: Electron transport rate, ΦPSII: Quantum efficiency of photosystem II in light, g_s_: Stomatal conductance, E: Transpiration rate, A_n_: Net CO_2_ assimilation rate and WUE_intr_cal_: Intrinsic water use efficiency. Square-root transformation was applied to WQ, g_s_ and E and log transformation (base 10) was applied to WUE_intr_cal_ before adjusting the linear mixed-model, as the data were not normally distributed.

There was a low but significant correlation between environments for leaf traits LMA, WC_f_, WQ and SPAD with r > 0.25 (Figure 2a diagonal). In contrast, no significant correlation was found between environments for ETR, ΦPSII, g_s_, E, A_n_ and WUE_intr_cal_.

The PCA performed on BLUPs from both greenhouse and field environments together (Figure 3a) revealed that the first component was positively associated with gas-exchange related traits and explained 22.4% of genotypic variability. Cultivars with positive coordinates on this first component had high values of A_n_ and g_s_ but low values of WUE_intr_cal_ whatever the environment. However, the second component, which explained 21.3% of the variability, showed contrasting patterns between environments, with traits contributing positively in one environment and negatively in the other. This was consistent with the absence of correlation between the two environments for gas-exchange related traits (Figures 2a diagonal and 3a). The third component was mainly explained by LMA and SPAD variations and the fourth one by the variation of WC_f_ and WQ. No discrimination of the environments was observed on these components (Figure 3b), in line with the correlations previously observed for these traits in the two environments (Figure 2a diagonal). PCA coordinates for each cultivar were also used in GWAS analyses, considering them as integrative traits.

**Figure 3:**
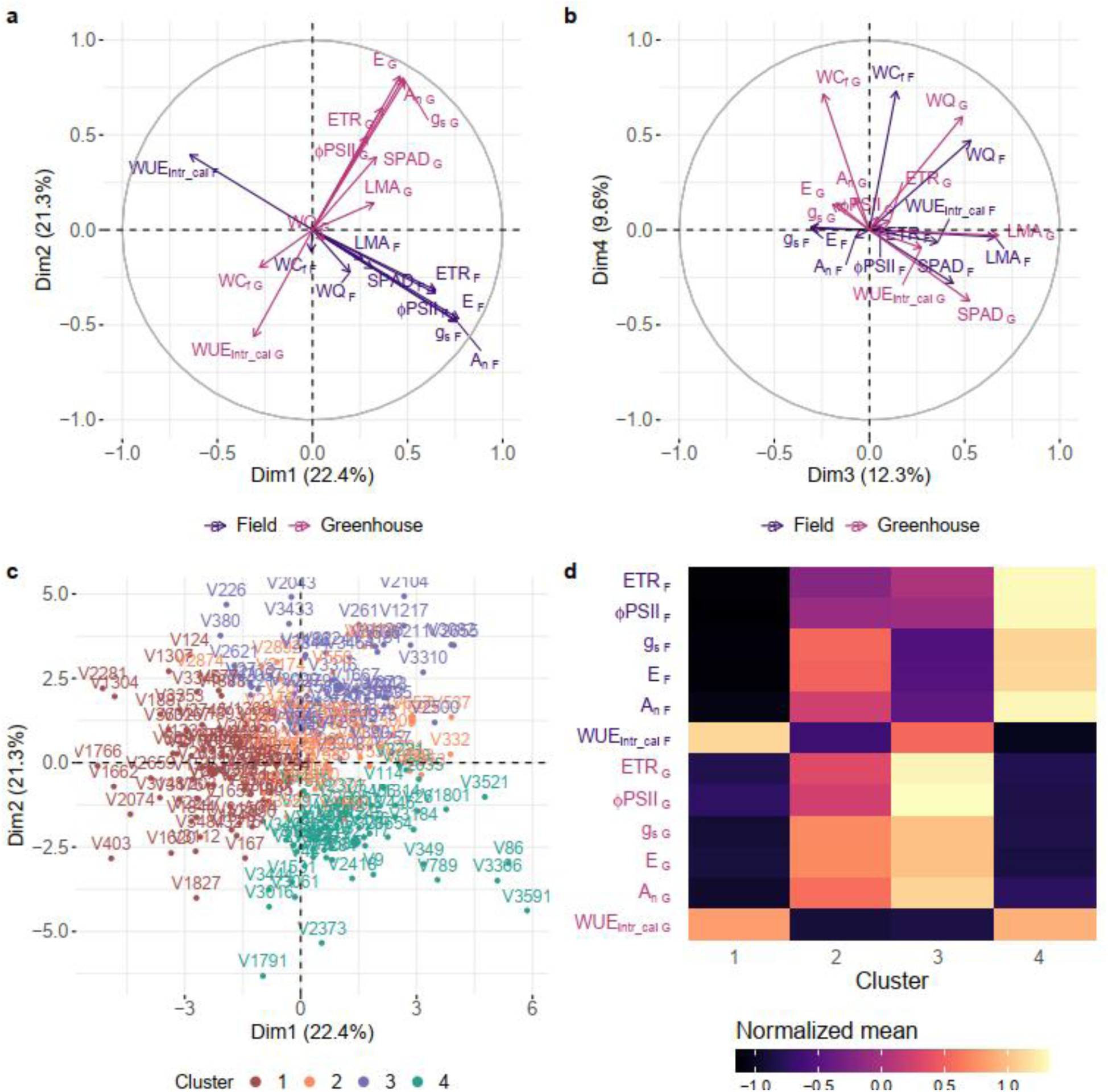
Multi-environment PCA and cultivar clustering. **a** Representation in the plane of the first two principal components of the PCA based on greenhouse (“G” subscript) and field (“F” subscript) trait BLUPs taken together. **b** Representation in the plane of the third and fourth principal components of the PCA based on greenhouse and field trait BLUPs. **c** Cultivars projection on the 1^st^ and 2^nd^ principal components of the same PCA, colored according to clusters from hierarchical clustering. Cultivars are designated with V followed by their IDVar, a unique cultivar identifier (freely available in Vassal-Montpellier Grapevine Biological Resources Center (Grapevine-BRC)). **d** Normalized means (centered and reduced) of BLUPs from both environments within each cluster, focusing on gas exchange-related traits. Cluster 1: 71 cultivars, Cluster 2: 44 cultivars, Cluster 3: 52 cultivars and Cluster 4: 54 cultivars. All analyses were performed on genotypic values (BLUPs) for n = 221 cultivars. LMA: Leaf mass per area, SPAD: Proxy of chlorophyll content, WC_f_: Water content, WQ: Water quantity, ETR: Electron transport rate, ΦPSII: Quantum efficiency of photosystem II in light, g_s_: Stomatal conductance, E: Transpiration rate, A_n_: Net CO_2_ assimilation rate and WUE_intr_cal_: Intrinsic water use efficiency. Square-root transformation was applied to WQ, g_s_ and E and log transformation (base 10) was applied to WUE_intr_cal_ before adjusting the linear mixed-model, as the data were not normally distributed.

Hierarchical clustering on principal components for the PCA combining greenhouse and field data, revealed four clusters of cultivars. They were mainly separated along the first two components, which reflected variations in gas exchange-related traits (Figure 3c). Cluster 1 included 71 cultivars with high values of WUE_intr_cal_ but low values of ETR, ΦPSII, g_s_, E and A_n_ whatever the environment (Figure 3d), whereas cluster 2 included 44 cultivars with low values of WUE_intr_cal_ but medium values of ETR and ΦPSII and high values of g_s_, E and A_n_ in both environments. Cultivars of clusters 3 (52 cultivars) and 4 (54 cultivars) were characterized by high trait values in the greenhouse and field environments, respectively, but an opposite pattern for WUE_intr_cal_.

### Genetic determinism of carbon- and water-related leaf traits

We used the genomic relationship matrix estimated from genome-wide markers to estimate the genomic heritability (h²_genomic_) of each trait in each environment. In general, h²_genomic_ in the field, ranging from 0.18 to 0.64, was often higher or equal to that in the greenhouse, where it ranged from 0.20 to 0.38 (Table 1). The highest h²_genomic_ were observed for LMA, WC_f_, WQ, SPAD and A_n_ in the field. These ranged from 0.46 to 0.64, nearly twice as high as in the greenhouse environment. h²_genomic_ of ETR and WUE_intr_cal_ were moderate, ranging from 0.18 to 0.38, with similar values for both environments. g_s_ and E also presented moderate h²_genomic_ values, but slightly higher in the greenhouse than in the field.

**Table 1:** Genomic heritability of BLUP leaf traits in greenhouse and field environments. The genomic heritability was calculated as follows: 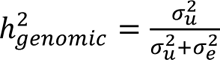 with *σ^2^_u_* the genetic variance and *σ*^2^ the residual variance, under the null model without individual variant effect. LMA: Leaf mass per area, SPAD: Proxy of chlorophyll content, WC_f_: Water content, WQ: Water quantity, ETR: Electron transport rate, ΦPSII: Quantum efficiency of photosystem II in light, g_s_: Stomatal conductance, E: Transpiration rate, A_n_: Net CO_2_ assimilation rate and WUE_intr_cal_: Intrinsic water use efficiency. Square-root transformation was applied to WQ, g_s_ and E and log transformation (base 10) was applied to WUE_intr_cal_ before adjusting the linear mixed-model, as the data were not normally distributed.

| Trait | h<sup>2</sup><sub>genomic</sub> Greenhouse | h<sup>2</sup><sub>genomic</sub> Field |
| --- | --- | --- |
| LMA | 0.32 | 0.58 |
| WC<sub>f</sub> | 0.25 | 0.52 |
| WQ | 0.35 | 0.64 |
| SPAD | 0.35 | 0.51 |
| ETR | 0.28 | 0.29 |
| ΦPSII | 0.28 | 0.41 |
| g<sub>s</sub> | 0.35 | 0.32 |
| E | 0.38 | 0.31 |
| A<sub>n</sub> | 0.26 | 0.46 |
| WUE<sub>intr_cal</sub> | 0.20 | 0.18 |

We further used different GWAS methods on the BLUPs, including single-trait *vs* multi-trait using PCA coordinates, single-*vs* multi-environment and single-locus mixed-model *vs* local score. QTL positions were defined as an interval around the detected variant based on the linkage disequilibrium (LD) decay for each chromosome (Table S5). A total of 132 QTLs controlling the leaf traits variability was identified (Table S6). Among those, 94 QTLs were found only in one environment. Six QTLs were detected using PCA coordinates performed on the data set combining greenhouse and field environment. Multi-environment GWAS methods (meta analyses (metaGE) and multi-trait mixed-model (MTMM)) allowed the detection of 32 additional QTLs: 27 QTLs with similar effects between both environments, primarily associated with ETR, ΦPSII, WUE_intr_cal_ and A_n_, and 5 QTLs with opposite effects, mainly associated with LMA and SPAD. Some QTLs were only detected for one trait and one method while others controlled different traits or were detected using multiple methods. Several QTLs common to gas exchange-related traits, such as ETR, ΦPSII, g_s_, E, A_n_ and WUE_intr_cal_ were identified, while no co-localization with the other traits (LMA, WC_f_, WQ and SPAD) was observed. The percentage of variance explained (PVE) by each topVariant of additive QTLs detected by single-trait and single-environment methods, ranged from 6.1% to 13.6%, with A_n_, E, ETR, ΦPSII and WUE_intr_cal_ having some QTLs with a PVE higher than 10%. The PVE ranged from 6.6% to 12.1% for additive single-environment PCA component traits.

We retained ten QTLs of particular interest for further analyses, based on the following criteria: (1) significance above the Bonferroni threshold; or (2) putative QTLs (*P*-value<10^-6^ or Lindley score above 9) with similar effects between environments, detected by both meta-analyses and MTMM methods, with possible co-localization among traits; or (3) putative QTLs (*P*-value<10^-6^ or Lindley score above 9) with opposite effects between environments, with possible co-localization among traits (Table 2).

**Table 2:**
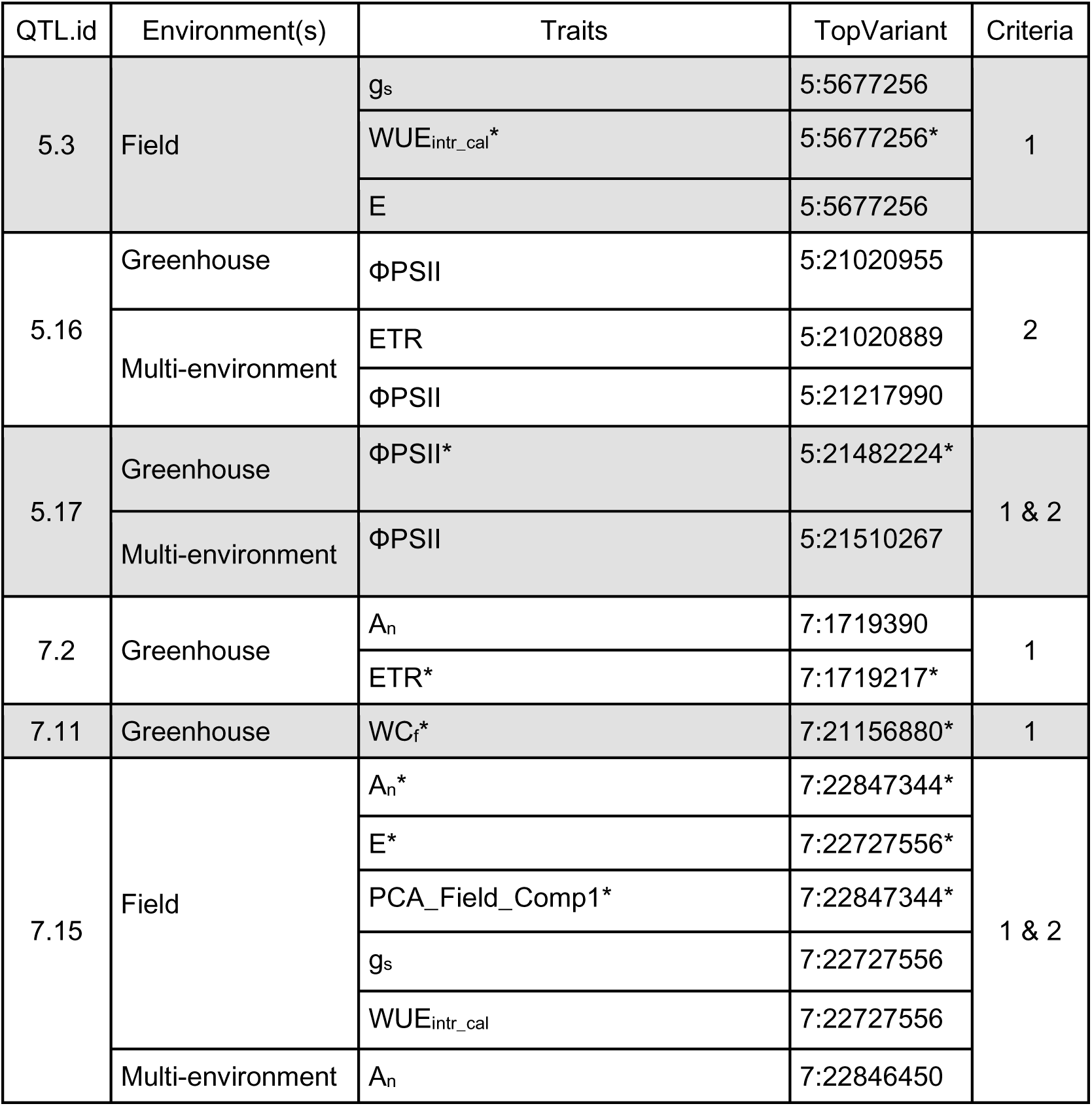
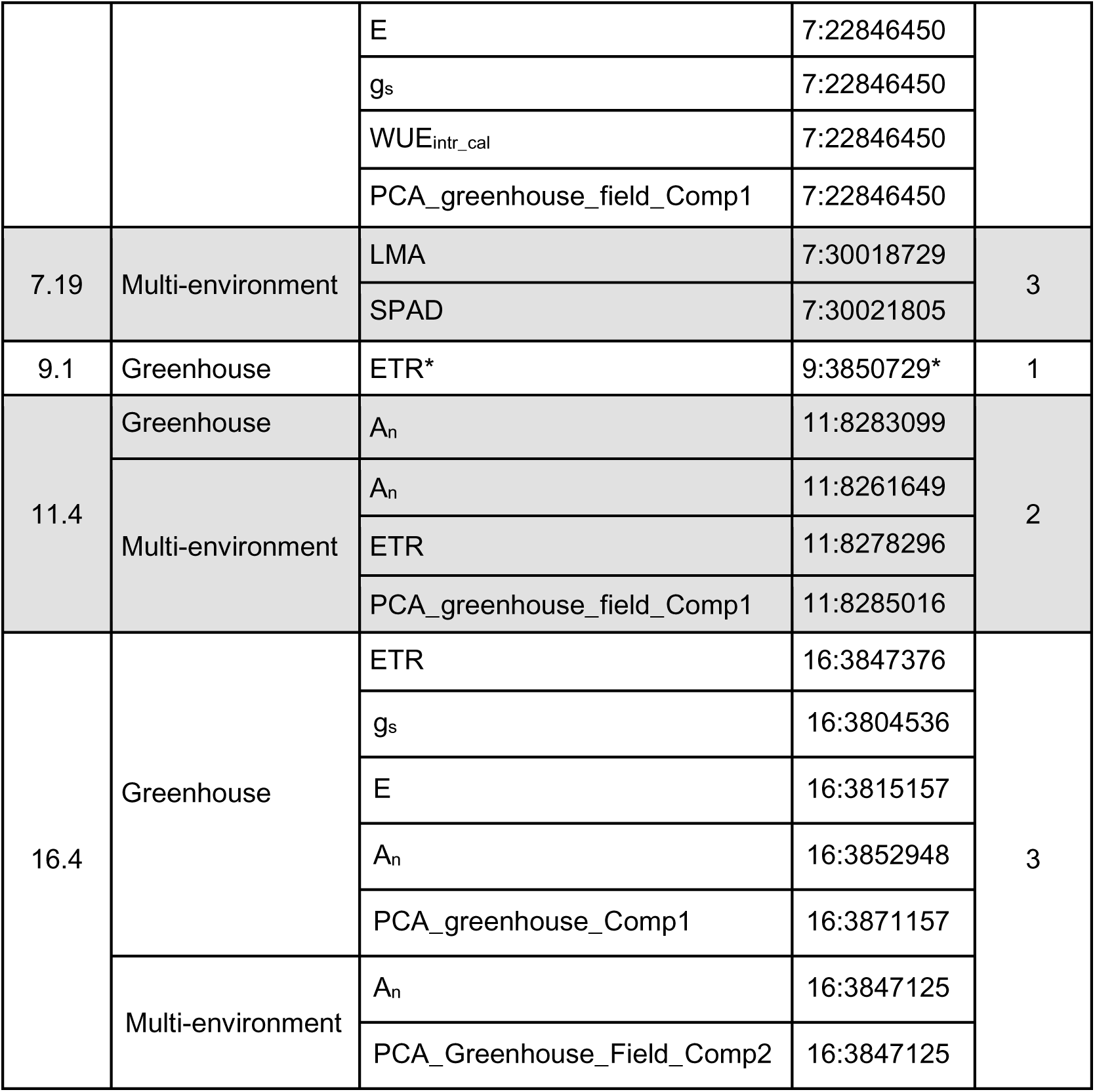
Summary of the subset of most interesting QTLs. This table is an extraction from Table S6 that describes all significant and putative QTLs found. A QTL was defined as an interval around the topVariant detected based on the linkage disequilibrium decay estimated for each chromosome. For overlapping QTLs, they were merged when LD between their topVariants was > 0.8. The QTLs presented here were retained if they met at least one of the following criteria: (1) significance above the Bonferroni threshold; or (2) putative QTLs (*P*-value<10^-6^ or Lindley score above 9) with similar effects between environments, detected by both meta-analyses and MTMM methods, with possible co-localization among traits; or (3) putative QTLs (*P*-value<10^-6^ or Lindley score above 9) with opposite effects between environments, with possible co-localization among traits. QTL.id: QTL identifier (composed of chromosome number and QTL order within the chromosome). Environment: the environment in which the QTL was found, either in greenhouse, in field or in both through multi-environment analyses. Traits: associated traits. TopVariant: identifier (chromosome:position in bp) of the most significant variant (either a SNP or a small indel) in the QTL for the given trait; * indicates topVariants with a -log_10_(*P*-value) greater than the Bonferroni threshold. Criteria: criteria (see above) that the QTL must satisfy for being retained in the subset. Some QTLs were detected with both single- and multi-environment models, thus they had two topVariants. LMA: Leaf mass per area, SPAD: Proxy of chlorophyll content, WC_f_: Water content, ETR: Electron transport rate, ΦPSII: Quantum efficiency of photosystem II in light, g_s_: Stomatal conductance, E: Transpiration rate, A_n_: Net CO_2_ assimilation rate and WUE_intr_cal_: Intrinsic water use efficiency, PCA_Greenhouse_Field_Comp1 (and PCA_Greenhouse_Field_Comp2): coordinate in the first (and second) principal component of the greenhouse+field PCA, PCA_Field_Comp1: coordinate in the first principal component of the field PCA and PCA_Greenhouse_Comp1: coordinate in the first principal component of the greenhouse PCA.

### QTLs detected in a single environment

Of all the QTLs found, 6 fulfilled the first criteria (significance above the Bonferroni threshold, 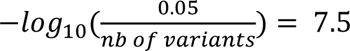 with a single-locus mixed-model on one environment either in field or greenhouse conditions (Table 2). Four of them were found in the greenhouse: QTLs 7.2 and 9.1 for ETR, 7.11 for WC_f_ and 5.17 for ΦPSII. The two other QTLs were found in the field: QTL 7.15 that controlled A_n_, E and the first principal component coordinate of Field PCA (PCA_Field_Comp1), and QTL 5.3 for WUE_intr_cal_. QTLs 5.17, 7.15 and 5.3 displayed an additive effect while QTLs 7.2, 9.1 and 7.11 showed a non-additive dominant or over-dominant effect (Figure S3, Figure 4).

**Figure 4:**
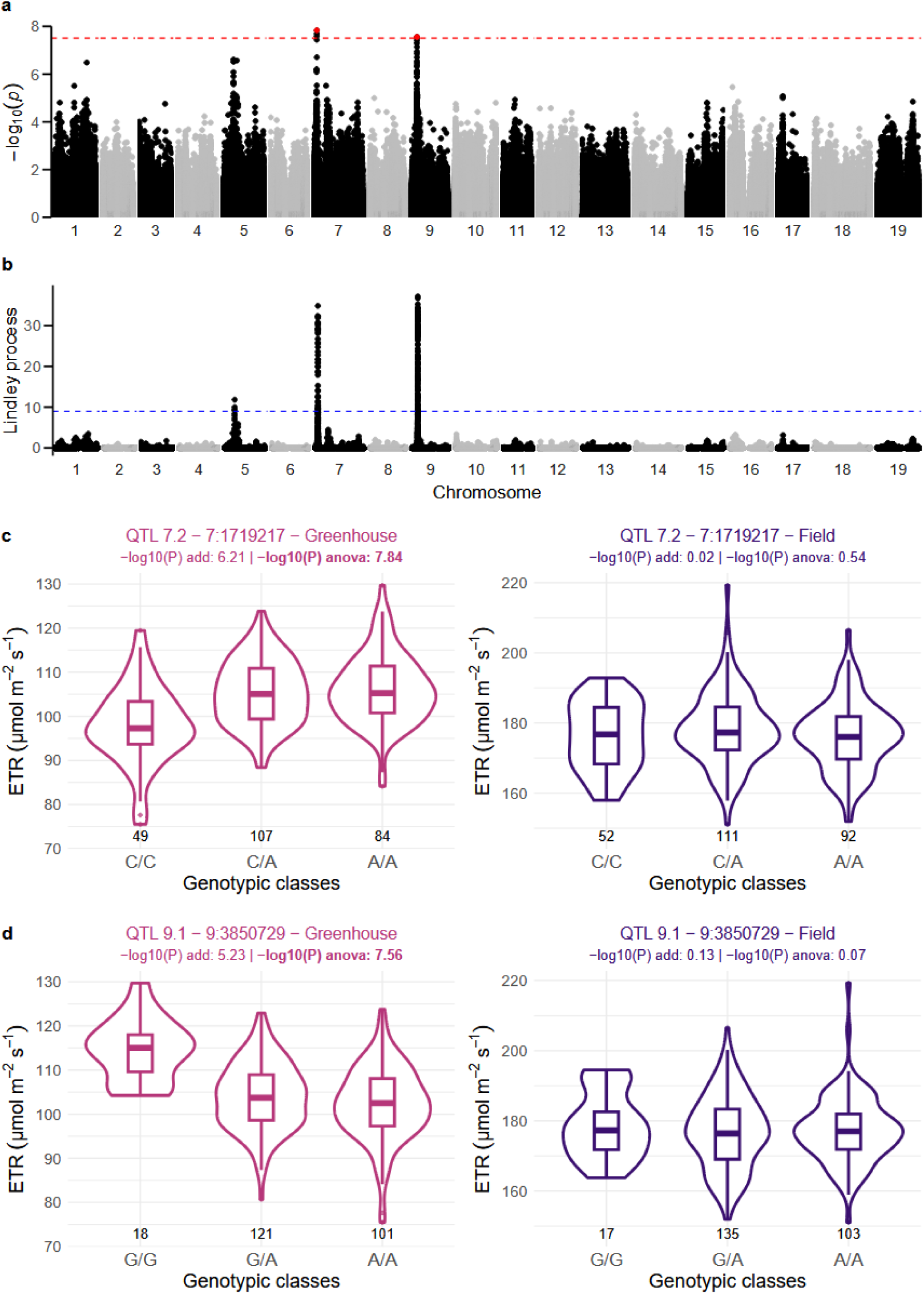
Detection of QTLs 7.2 and 9.1 via single-environment GWAS for genotypic values (BLUPs) of electron transport rate (ETR) using the single-locus MM4LMM method. **a** Genome-wide Manhattan plot for the greenhouse environment associations obtained with a single-locus mixed-model adapted to identify both additive and non-additive effects (ANOVA). Red dots are for 7:1719217 and 9:3850729 topSNPs of QTLs 7.2 and 9.1, respectively. The red dashed line corresponds to the Bonferroni threshold 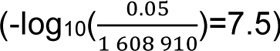. **b** Lindley process along chromosomes from Local Score approach based on the *P*-values from the single-locus mixed-model adapted to identify both additive and non-additive effects (shown in **a**). The blue dashed line is the Local Score threshold (Lindley score of 9). **c** and **d** Boxplots and violinplots showing the effect of the 7:1719217 topSNP genotypic classes of QTL 7.2 (**c**) and the effect of the 9:3850729 topSNP genotypic classes of QTL 9.1 (**d**) on ETR genotypic values (BLUPs) in the greenhouse and field environments (in µmol m^-2^ s^-1^). The number of cultivars with each genotypic class is given under the corresponding boxplot. -log_10_(P) add is the -log_10_(*P*-value) for the additive only model and -log_10_(P) anova is the -log_10_(*P*-value) for the additive + non-additive model. 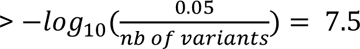 are highlighted in bold.

QTLs 7.2 and 9.1 were both significant at the Bonferroni threshold for ETR. Their topVariants were the SNPs 7:1719217 and 9:3850729 which explained 9% and 8% of the trait, respectively (Table S6). The Manhattan plot revealed a peak below these topSNPs, where neighbouring variants had -log_10_(*P*-value) greater than 3 and were in linkage disequilibrium with the associated topSNP. This raised the Lindley score for these topSNPs, computed with the local score approach, above 30 (Figure 4a and b). In the greenhouse, QTL 7.2 was not only associated with ETR but also with A_n_ (-log_10_(*P*-value) > 6). Its topSNP for ETR (7:1719217) had a dominant effect (Figure 4c), with cultivars carrying at least one alternate allele A (compared to the reference genome) having higher ETR and A_n_ than the other cultivars. The dominant effect observed for 9:3850729 resulted in lower ETR values for cultivars with at least one copy of the alternate allele A (Figure 4d). However, unlike QTL 7.2, this topSNP had no effect on A_n_ (–log_10_(*P*-value) < 5). None of these 2 topSNPs with dominant effects was associated with ETR in the field single-environment GWAS (-log_10_(*P*-value) < 0.54) (Figure 4c and d). We found relevant candidate genes in close proximity to these two topSNPs. QTL 7.2 contained the *Vitvi05_01chr07g02400* gene, homologous to a gene encoding a plastocyanin protein involved in the photosynthetic electron transport chain in *Arabidopsis thaliana* (Table S7). Under QTL 9.1, we found the *Vitvi05_01chr09g04720* gene homologous to a gene encoding a pentatricopeptide repeat (PPR)–containing protein required for NDH complex activity in the photosynthetic electron transport chain.

### QTLs with consistent effects across environments

None of the QTLs found using single-environment GWAS (either significant or putative) was common in both greenhouse and field (Table 2, Table S6). Hence, to specifically investigate whether some QTLs could be shared between environments, with effects either in the same or opposite directions, we examined multi-environment GWAS results from meta-analyses and MTMM methods. Among the QTLs thus detected, four were of particular interest, as they displayed similar shared effects across environments (Table 2): 5.16, 5.17, 7.15 and 11.4. QTLs. 5.17 and 7.15 had already been detected using either greenhouse or field single-environment GWAS, respectively (Criteria 1 Table 2, Figure S3 b, d-e). However, topVariants were not identical between single- and multi-environment GWAS (Figure S4 b-c).

For ΦPSII, QTLs 5.16 and 5.17 had an additive effect in both environments (Figure S4 a-b). The candidate gene *Vitvi05_01chr05g20200* was found in the interval underlying QTL 5.16 and appeared of particular relevance because it was homologous to YLMG1 in *Arabidopsis thaliana*. This gene is likely involved in chloroplast division, and YLMG1 silencing in *A. thaliana* led to pale green leaves which could negatively impact ΦPSII (Table S7).

QTL 7.15 was found for An, E, gs, WUEintr_cal, as well as for PCA_greenhouse_field_Comp1. We observed strong LD within this QTL (r²V > 0.6) between the topVariant 7:22846450 and some upstream variants (Figure S5a) where six relevant candidate genes were identified based on their gene annotation (Table S7). Three of them were homologous to *A. thaliana* genes involved in adaxial-abaxial leaf polarity development (*Vitvi05_01chr07g22140*, *Vitvi05_01chr07g22160* and *Vitvi05_01chr07g22170*) and three in the control of stomatal opening or closure (*Vitvi05_01chr07g22080* annotated as a Serine/threonine-protein kinase, *Vitvi05_01chr07g22250* and *Vitvi05_01chr07g22260* annotated as a phospholipase PLA2). *Vitvi05_01chr07g22260*, *Vitvi05_01chr07g22140* and *Vitvi05_01chr07g22080* have been reported to be frequently expressed in grapevine leaves (Figure S5b, Table S7).

QTL 11.4 was detected for A_n_, ETR and PCA_greenhouse_field_Comp1. Its topSNP 11:8261649 displayed an additive effect for A_n_ in both greenhouse and field environments with cultivars homozygous for the reference allele C having lower A_n_ compared to cultivars homozygous for the alternate allele T (Figure 5c and d). Of note, LD was rather weak around the topSNP (Figure 5a). However, within the QTL region 336 variants had a -log_10_(*P*-value) > 3 and showed consistent allelic effects, which positively contributed to the Lindley score that reached a maximum value of 81 (Table S6). A *χ*² test on the distribution of cultivars per cluster from the greenhouse+field PCA (Figure 3, Figure 5e) showed an enrichment in cultivars from cluster 1 (with leaves presenting low A_n_ regardless of the environment, Figure 3d) in the reference allele homozygous genotypic class (C/C). Gene annotation in the QTL further highlighted 4 candidate genes of interest: *Vitvi05_01chr11g10240*, *Vitvi05_01chr11g10280*and *Vitvi05_01chr11g10300*, *Vitvi05_01chr11g10310*, homologous to genes that, in *A. thaliana*, either exhibited a photosynthetic defective mutants with a pale phenotype or participated in electron transfer (Table S7). These four genes were reported to be frequently highly expressed in grapevine leaves (Figure 5b).

**Figure 5:**
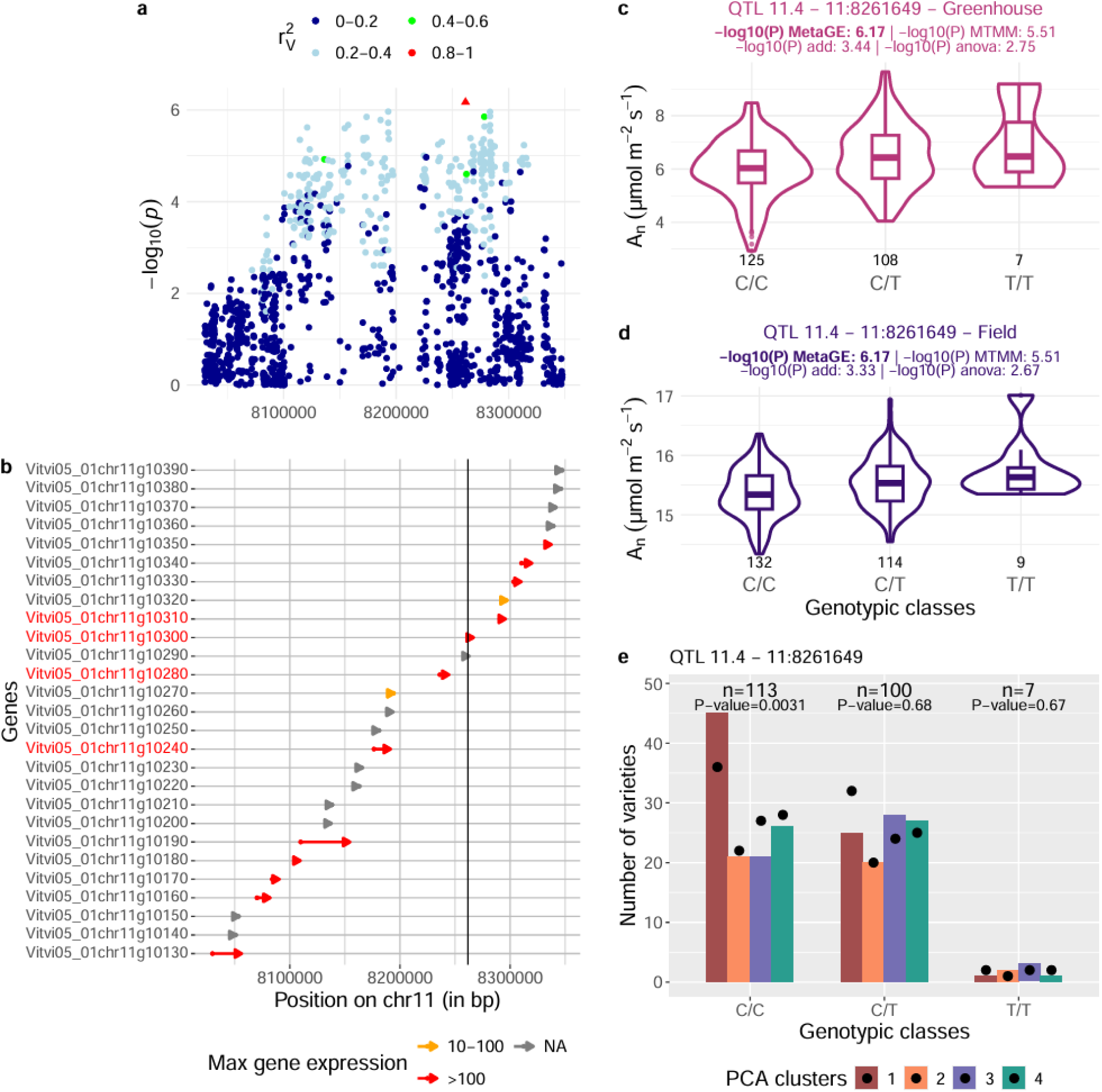
Fine characterisation of the QTL 11.4 having a similar effect for genotypic values (BLUPs) of net CO2 assimilation rate (An) between both greenhouse and field environments, detected by the meta analysis and the common approach from the MTMM method, combined with local score approach. **a** Zoom in the genome-wide Manhattan plot performed using a meta analysis for A_n_ in both greenhouse and field environments. Colors correspond to the local linkage disequilibrium corrected by the relatedness between cultivars (r²_V_) based on the topSNP 11:8261649 (triangle) of the QTL 11.4. **b** Maximum gene expression in leaves (color of the arrows) extracted from GREAT (GRape Expression Atlas) RNA-Seq data analysis workflow for the “leaf” organ across all studies, expressed in Reads Per Kilobase of transcript per Million reads mapped (RPKM). The black line corresponds to the position of the topSNP, aligned with **a**. Genes with red names are those with a putative function related to the trait of interest. The size of the arrow is proportional to the size of the gene. **c** and **d** Boxplots and violinplots showing the effect of the 11:8261649 topSNP genotypic classes of QTL 11.4 on A_n_ for the greenhouse and field environments, respectively (in µmol m^-2^ s^-1^). The number of cultivars with each genotypic class is given under the corresponding boxplot. -log_10_(P) MetaGE is the -log_10_(*P*-value) for the meta-analysis model and -log_10_(P) MTMM is the -log_10_(*P*-value) for the MTMM common approach. -log_10_(P) add is the -log_10_(*P*-value) for the additive only model and -log_10_(P) anova is the -log_10_(*P*-value) for the additive + non-additive model using the single-environment single-locus mixed-model. *P*-value<10^-6^ are highlighted in bold. **e** Barplot of the distribution of cultivars according to the four PCA clusters for the three genotypic classes of 11:8261649. The black dot is the expected number of cultivars in each cluster for each genotypic class under the null hypothesis of no association between genotypic class and PCA cluster. The *P*-value of the *χ*² test for each genotypic class is above the bars.

### QTLs with opposite effects across environments

Finally, we specifically examined two QTLs with opposite effects across environments with the gei effect test of the MTMM model (Criteria 3, Table 2). One of those was QTL 7.19, found for LMA (Figure S4d). Another was QTL 16.4, associated with the variations in ETR, A_n_, g_s_ and E. This QTL was previously identified by the single-environment single-locus mixed-model in the greenhouse as well as for PCA_greenhouse_Comp1 (Table 2, Figure 2c). The opposite effect between environments of this QTL was identified through the gei model of MTMM method and supported by its detection for PCA_Greenhouse_Field_Comp2, the second component from the PCA in which traits variations were orthogonal between environments (Figure 3a).

For this QTL, there were 42 variants in high LD (R²_V_ >0.8, Figure 6a) with a -log_10_(*P*-value) higher than 3 in the region of the QTL 16.4, which led to a strong signal detection through the Local Score approach on the result of gei model of MTMM method (Figure S6), with the identification of the topSNP 16:3847125. This topSNP displayed opposite additive effects on A_n_ between environments, with the reference allele (A) being associated with higher A_n_ values in the greenhouse (Figure 6c) and lower A_n_ values in the field, compared with the alternate allele (Figure 6d). Consistently, opposite effects between environments were also found for ETR, g_s_ and E (data not shown). Moreover, a significant disequilibrium in the distribution of cultivars across PCA clusters was observed for each genotypic class (Figure 6e). The number of cultivars in the reference allele homozygous class (A/A) was half that expected for cluster 4, where cultivars had higher values of ETR, ΦPSII, g_s_, E, and A_n_ in the field than in the greenhouse (Figure 3d). On the contrary, there were significantly more cultivars from cluster 4 with the alternate allele which increased the traits values in the field. *Vitvi05_01chr16g04040*, *Vitvi05_01chr16g04070*, *Vitvi05_01chr16g04130* and *Vitvi05_01chr16g04140* genes were found as relevant candidate genes in this QTL 16.4. Three out of these four genes were reported to be expressed in grapevine leaves (Figure 6b). They were homologous to genes in *A. thaliana* encoding for pectin methylesterase proteins which play an important role in stomatal development and functioning (Figure 6b, Table S7).

**Figure 6:**
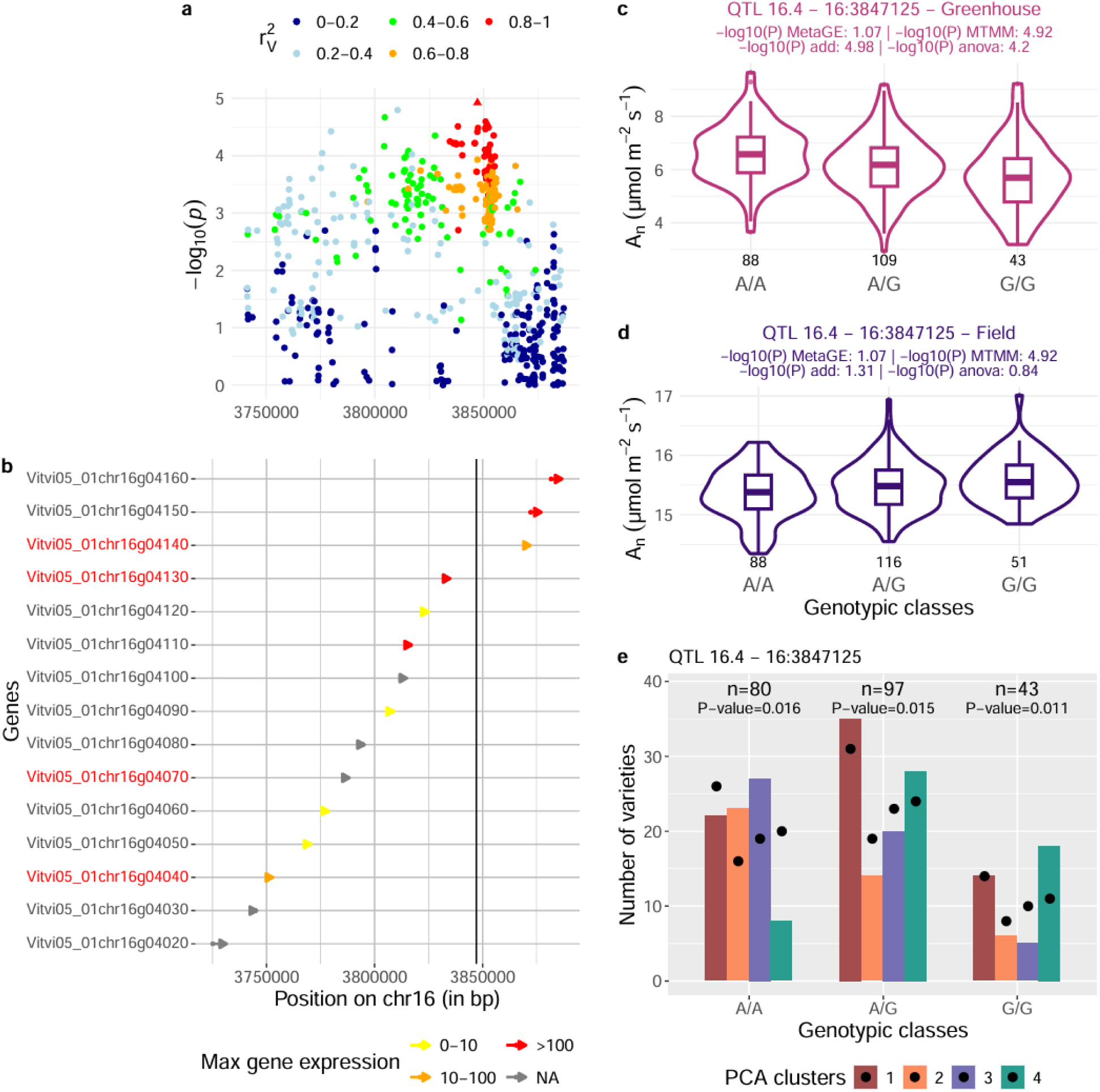
Fine characterisation of the QTL 16.4 with an interaction effect for genotypic values (BLUPs) of net CO2 assimilation rate (An) between greenhouse and field environments, detected by gei model of MTMM method combined with local score approach. **a** Zoom in the genome-wide Manhattan plot performed using a single-locus multi-environment mixed-model for A_n_ from both greenhouse and field environments. Colors correspond to the local linkage disequilibrium corrected by the relatedness between cultivars (r²_V_) based on the topSNP 16:3847125 (triangle) of the QTL 16.4. **b** Maximum gene expression in leaves (color of the arrows) extracted from GREAT (Grape Expression Atlas) RNA-Seq data analysis workflow for the “leaf” organ across all studies, expressed in Reads Per Kilobase of transcript per Million reads mapped (RPKM). The black line corresponds to the position of the topSNP, aligned with **a**. Genes with red names are those with a putative function related to the trait of interest. The size of the arrow is proportional to the size of the gene. **c** and **d** Boxplots and violinplots showing the effect of the 16:3847125 topSNP genotypic classes of QTL 16.4 on A_n_ for the greenhouse and field environments, respectively (in µmol m^-2^ s^-1^). The number of cultivars with each genotypic class is given under the corresponding boxplot. -log_10_(P) MetaGE is the -log_10_(*P*-value) for the meta-analysis model and -log_10_(P) MTMM is the -log_10_(*P*-value) for the MTMM approach. -log_10_(P) add is the -log_10_(*P*-value) for the additive only model and -log_10_(P) anova is the -log_10_(*P*-value) for the additive + non-additive model using the single-environment single-locus mixed-model. **e** Barplot of the distribution of cultivars according to the four PCA clusters for the three genotypic classes of 16:3847125. The black dot is the expected number of cultivars in each cluster for each genotypic class under the null hypothesis of no association between genotypic class and PCA cluster. The *P*-value of the *χ*² test for each genotypic class is above the bars.

## Discussion

The main objective of this study was to assess how the genetic variability and determinism of carbon- and water-related leaf traits in grapevine are conserved between environments (greenhouse and field conditions respectively). We deployed an original combination of high-throughput leaf phenotyping, dense genotyping and complementary GWAS methods to address this question on a large diversity panel of grapevine. We first observed that, although the traits showed similar correlation patterns within each environment, the same traits measured in greenhouse and field environments were poorly correlated. Then, we characterized the genetic determinism of these traits and found QTLs with specific effects within each environment using single-environment GWAS. Finally, we carried out multi-environment GWAS, which highlighted QTLs with shared effects across both environments. We highlighted relevant candidate genes within the QTLs to help identify the underlying physiological processes.

### Comparison greenhouse vs field: a robust experimental and analytical strategy but major management differences

We measured in each environment more than a thousand leaves for a number of functioning and morphological traits related to carbon and water. As expected, environmental effects such as date and time of measurements, and micro climatic variables, significantly impacted the observed variability. Yet, the adequate experimental design combined with the use of mixed models allowed us to account for an important part of this undesired variability to ensure the accurate estimation of genotypic values (BLUPs). Based on these reliable BLUPs we identified stable QTLs between both environments. The robustness of these QTLs was also supported by the identification of candidate genes relevant to the traits, such as *Vitvi05_01chr07g22260* and *Vitvi05_01chr07g22080* which are involved in the regulation of stomatal opening and were present in QTL 7.15, identified in particular for E and g_s_.

It is important to note, however, that several specificities in plant management strongly differentiated both experiments. One major difference was the growing conditions: plants were grown in pots in the greenhouse, while they were grown directly in the soil in the field. On the one hand, the use of potting soil in potted experiments, with high water retention capacity and low resistance to root growth (Poorter et al., 2016) allows to reduce the spatial variability and soil heterogeneity often observed under field conditions. However, on the other hand, pot size restricts root development, which may limit the expression of genotypic differences in root growth across cultivars. In addition, pot size may also increase soil and root temperatures, potentially affecting root growth (Poorter et al., 2012) and influence water and nutrient uptake, which may lead to hypoxic conditions that negatively affect the root system (Passioura, 2006). These constraints may ultimately affect plant growth, including through reduced photosynthetic activity (Poorter et al., 2012). Such effects could partly explain the lower values of A_n_ observed here in the greenhouse compared with those observed in the field.

The comparison of genotypic values between environments may also be hindered by differences in grafting conditions. In the greenhouse experiment, plants were grown on their own roots whereas they were grafted on SO4 rootstock (*Vitis berlandieri* x *Vitis riparia*) in the field. Significant rootstock effect, as well as scion x rootstock interaction effect, have been reported not only on plant total dry weight (Tandonnet et al., 2009) but also on aerial functioning including transpiration (Marguerit et al., 2012), stomatal conductance and photosynthesis (Zhang et al., 2016). The overall higher values in gas-exchange related traits as well as the changes in cultivars ranking for these traits that we observed in the field experiment as compared to the greenhouse conditions could also arise from these effects. Despite those differences, we identified a number of QTLs with a similar and consistent effect across both environments (for ETR, ΦPSII, A_n_, E, g_s_ and WUE_intr_cal_). This suggests that some of the genetic determinants of ecophysiological traits are relatively robust to differences in root systems.

Finally, the mean air temperature was 31.6 and 26.2°C in the greenhouse and in the field, respectively. Within this range of temperature the photosynthetically photon flux density can be considered saturating at a minimum of 800 µmol m^-2^ s^-1^ (Greer & Weedon, 2012). In this case, more than 50% of the measurements were carried out under non-saturating light in the greenhouse compared with 10% in the field. This difference in light conditions may explain the observed differences in A_n_, as well as leaf morphological traits, with plants grown in the greenhouse exhibiting thinner leaves (lower LMA values) but higher chlorophyll content per unit leaf area (SPAD) than field-grown plants. These results corroborate a previous wheat study where plants under greenhouse conditions exhibited significantly higher nitrogen and chlorophyll contents than in field conditions (Sales et al., 2022).

### Added value of the combination of different genome wide analyses methods to detect QTLs

The use of appropriate statistical methods was crucial, both because of the weak inter-environment correlation for each physiological trait and also because of the complex, polygenic nature of the traits. Despite the major management differences between experiments, multi-environment approaches (MTMM model and meta-analysis) allowed to identify up to 4 QTLs with consistent effect in both environments out of the 10 QTLs of interest. They therefore provide a valuable complement to single-environment approaches.

To adequately handle the new, very dense genotyping matrix used, we chose to use the local score approach recently proposed (Bonhomme et al., 2019; Fariello et al., 2017). In such GWAS settings, the main difficulty is to detect significant variants when a considerable number of variants have small effects (Barton et al., 2017). This method addresses the problem by accumulating association signals across consecutive variants according to their genomic location and can therefore detect small-effect QTLs explaining 5 to 10% of the genotypic variation. In our study, 8 of the 10 QTLs of interest were detected using the local score approach, compared to 6 based on the Bonferroni threshold. Notably, 4 of the 8 QTLs detected by the local score approach did not reach the Bonferroni threshold, highlighting its ability to capture cumulative moderate association signals rather than one single highly significant variant. The QTL 11.4 particularly illustrates the added value of using this approach as its Lindley score reached 80 whereas the most strongly associated SNP in this region had a *P-*value just under 10^-6^. Thus, this QTL would have remained undetected using a conventional single-locus mixed-model approach using a Bonferroni threshold, whereas the local score approach was able to capture the combined contribution of multiple modest association signals within the genomic region.

In addition to improving the detection of QTLs with small effects, another originality of this work was to investigate non-additive associations between variants and traits. This aspect is particularly relevant in grapevine, as more than 30% of its genome is heterozygous (Laucou et al., 2018). In previous studies, additive and dominance variances were influenced by mean heterozygosity: additive variance decreased for medium to high heterozygosity, while dominant variance increased (Nietlisbach et al., 2016), yielding the detection of non-additive QTLs. Recently, studies on *Prunus* species have highlighted the significant role of non-additive QTLs in controlling agronomic traits (Pérez De Los Cobos et al., 2023) and biotic stress responses (Serrie et al., 2025). In this study, 3 of the 10 QTLs of interest showed non-additive associations and would not necessarily have been detected using an additive single-environment GWAS model. However, we were unable to investigate non-additive effects in multi-environment models as these have not yet been implemented to detect non-additive effects.

### QTLs classification according to two physiological processes

Despite the correlations observed between carbon- and water-related traits (particularly A_n_, ETR, ΦPSII, E and g_s_), which were consistent across both environments, and the clustering of cultivars according to traits and/or environments, our results suggest that the identified QTLs can be broadly grouped according to two distinct physiological processes. These processes appear to be related to the stomatal control and the functioning of the photosynthetic apparatus, respectively.

QTLs such as 7.15 have been identified for both carbon-(A_n_) and water-related traits (E and g_s_), as well as for WUE_intr_cal_ and the coordinate of the first principal component of PCA from greenhouse+field PCA which discriminated between these traits. Moreover, the QTL 7.15 has already been reported for g_s_ in the same grapevine diversity panel using a lower density matrix of 80k SNPs (Chir et al., 2025). Some candidate genes identified within this QTL were annotated as phospholipase PLA_2_, involved in the control of stomatal opening in response to light (Seo et al., 2008). This raises the possibility that the effect of this QTL on A_n_ is mediated indirectly through its regulation of stomatal opening.

On the other hand, some QTLs can be grouped as potentially involved in the photosynthetic machinery. This is the case for QTLs 5.16 and 11.4, which have been detected for carbon-related traits such as ETR and ΦPSII or A_n_. The *A. thaliana* homologs of the candidate genes underlying these QTLs: YLMG1-1 homolog of *Vitvi05_01chr05g20200* in QTL 5.16, LPAT1 homolog of *Vitvi05_01chr11g10240* in QTL 11.4, were associated with a pale-green leaf phenotype and significant photosynthesis defect in their mutants (Barroga et al., 2024; Kabeya et al., 2010; Kobayashi et al., 2007). This suggests that these genes may be involved in the functioning of the photosynthetic apparatus.

Overall, our results suggest the existence of stable genetic determinants of leaf functioning traits, as several QTLs were stable across environments. Moreover, the genetic determinants associated with water loss and carbon assimilation appear to be at least partly distinct. This provides promising insights for breeding strategies aiming at maintaining net carbon assimilation without necessarily increasing water loss, thereby contributing to the improvement of cultivars under water deficit conditions.

## Materials and methods

### Plant material and growing conditions

The diversity panel consisted of 279 grapevine cultivars. This panel was designed to capture the genetic diversity of *V. vinifera* (Nicolas et al., 2016). It was studied in two distinct experiments: one on potted plants hosted in a greenhouse, and one in the field.

The experiment carried out in the greenhouse, previously described in Coindre et al., 2026, was conducted in 2022 using 241 cultivars from the diversity panel. Six four-years-old own-rooted plants per cultivar were grown in pots and phenotyped in the PhenoArch phenotyping platform (Cabrera-Bosquet et al., 2016) hosted at the M3P (Montpellier Plant Phenotyping Platforms, https://eng-lepse.montpellier.hub.inrae.fr/platforms-m3p/montpellier-plant-phenotyping-platforms-m3p). Three different watering treatments were set up, corresponding to the following target soil water potentials: −0.03 MPa for well-watered condition (WW), −0.11 MPa for moderate water deficit (WD1) and −0.43 MPa for severe water deficit (WD2). Most genotypes included two replicates (i.e. two plants) in each water treatment (six plants in total). Leaf phenotyping was conducted over a total of 24 mornings during June and July 2022. Growing conditions and water treatment management were previously described (Coindre et al., 2026). Ambient

Photosynthetically Photon Flux Density (PPFD) was monitored with SKP 215 (Skye Instruments), air temperature (T_air_) and relative humidity were monitored with a Rotronic HC2-SH (Rotronic AG), enabling the calculation of Vapor Pressure Deficit (VPD). These probes were distributed throughout 10 different locations in the greenhouse.

The experiment conducted in the field was performed in 2024. It included 255 cultivars of the diversity panel and shared 222 common cultivars with the greenhouse experiment. All the vines were grafted on SO4 rootstock (*Vitis berlandieri* x *Vitis riparia*) and planted in the field in 2021 in four randomized complete blocks at Pech Rouge INRAE experimental vineyard, Gruissan (France). Each block was divided into unit plots of five contiguous plants of the same cultivar. Each plant was drip-irrigated the night before phenotyping. Leaf phenotyping was conducted on two blocks over 4 sunny mornings in June 2024.

PPFD, T_air_ and VPD were recorded at the leaf level with a hand-held poro-fluorometer (Li600, Portable Photosynthesis System LI-600, Licor, Lincoln, United States) at the time of measurements in the field experiment.

### Phenotyping or prediction of leaf traits

The leaf phenotyping was performed between 9:00 and 13:30 (UTC+2) for both greenhouse and field experiments. In the greenhouse, we measured one leaf per plant for each of the six plants of each cultivar. In the field, we measured two leaves per cultivar per block for two blocks, over two of the three central plants of the unit plot of five plants (one leaf per plant). In both experiments, fully expanded leaves were chosen on primary shoots, well exposed to sunlight and usually located between the tenth and the fifteenth phytomer from the apex. A total of 1,449 and 1,102 leaves were phenotyped in the greenhouse and field experiments, respectively (Table S1).

The phenotyping protocol was described in (Coindre et al., 2026) and is detailed here in Supplementary Method 1. Briefly, transpiration rate (E, mmol m^-2^ s^-1^), stomatal conductance (g_s_, mol m^-2^ s^-1^), quantum efficiency of photosystem II in light (ΦPSII) and electron transport rate (ETR, µmol m^-2^ s^-1^) were measured using a hand-held poro-fluorometer (Li600, Portable Photosynthesis System LI-600, Licor, Lincoln, United States). A proxy of chlorophyll content was assessed as SPAD value using SPAD-502 Chlorophyll Meter (Konica-Minolta, Tokyo, Japan). Some traits could not be directly measured on all leaves. Instead, they were derived from high-throughput measurements, measured with spectrometers (MicroNIR™ Onsite-W, VIAVI, Chandler, AZ, United States or ASD LabSpec® 4, Analytical Spectral Devices, Boulder, USA) or poro-fluorometer (Li600), using prediction models built with the Partial Least Square Regression (PLSR) method, as implemented in R/pls (Liland et al., 1999). Thus, Leaf Mass per Area (LMA, mg cm^-2^), Water Quantity (WQ, mg H_2_O per leaf disk) and Water Content (WCf, mg H_2_O mg^-1^ fresh weight) were directly measured in the greenhouse environment but predicted in the field (Table S1, Table S2) while, Net CO_2_ Assimilation rate (A_n_, µmol m^-2^ s^-1^) was predicted for the greenhouse (Coindre et al., 2026) and the field experiment (for more details on trait prediction using PLSR models, see Supplementary Method 2). Intrinsic Water Use Efficiency (WUE_intr_cal_, µmol CO_2_ mol^-1^ H_2_O) was not well predicted in the greenhouse experiment based on PLSR model so we rather chose to calculate it in both experiments by dividing the predicted values of A_n_ by the measured values of g_s_ obtained with the Li600 device (Figure S7a). It was calculated for all plants except for plants under a severe water deficit where the predictive quality was limited (Figure S7b).

In the end, each leaf from each plant in the greenhouse and field experiments had a value for each trait, either through measurement or prediction (Table S1). All data analyses were handled in R v4.3.1 (R Core Team, 2023).

### Linear mixed-model and estimation of genotypic values

Genotypic values, corrected from experimental design and climatic effects, were extracted for each trait (either directly measured or predicted through PLSR) by fitting a linear mixed-model using R/lme4 (Bates et al., 2015) for each environment, as follows:

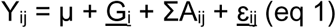

where Y_ij_ was the phenotypic trait value measured on plant j (j from 1 to 6) from cultivar i (i from 1 to 255), µ was the overall mean, G_i_ was the random effect of cultivar i, ΣA_ij_ was the sum of fixed effects of experimental covariates and ɛ_ij_ was the random residual effect. For the greenhouse environment, the interaction between cultivar and water treatment was also included into the model as a random effect. The date and time of measurement were included as fixed effects in both environments. In the greenhouse environment, additional fixed effects included water treatment, the time elapsed between the last irrigation of the plant and the measurement, the duration in days between the entrance of the plant into the greenhouse and the measurement, the plant density around each plant and the mean T_air_, PPFD and VPD between the entrance of the plant into the greenhouse and the measurement (recorded by probes). In the field environment, fixed effects included the block, the row and the position of the plot in the row considered as factors as well as T_air_, PPFD, VPD (all three recorded by the Li600 device) and the hourly wind speed (recorded by the weather station ca. 100 m from the field). In all cases, a model selection was performed for each trait to retain only significant fixed effects (*P*-value < 0.05), using plantTrialLmmFitCompSel from R/rutilstimflutre (Flutre & Chabrault, 2019). Genotypic Best Linear Unbiased Predictors (BLUPs) were extracted from these models and subsequently used as environment-specific estimates of genetic values (Table S4). For both environments, as g_s_, E and WQ were not symmetrically distributed, square-root transformation was applied to these traits values before adjusting the linear mixed-model while WUE_intr_cal_ was log-transformed (base 10) (Table S4).

### Principal component analysis and hierarchical clustering

In order to study relationships between traits, Pearson correlation coefficients between genotypic BLUPs were calculated and Principal Component Analyses (PCA) were carried out using R/FactoMineR (Husson, 2010), either considering each environment separately or both environments together. Principal component axes explaining more than 10% of the variability were then used as additional, synthetic traits for genetic analyses.

A hierarchical ascendant clustering on principal components (HCPC), using Ward’s agglomerative method based on Euclidean distances between PCA scores, was applied to the PCA combining the two environments to identify clusters of cultivars with similar leaf functioning properties. The number of clusters was selected from the dendrogram, using the main break in the hierarchy in linkage distance as the criterion. In this study, the number of clusters was set to 4.

### Genotyping data

Variant genotyping was performed using whole genome short-read resequencing data aligned on the *V. vinifera* PN40024_T2T (hereafter referred to as PN_T2T) v5 reference genome assembly (Shi et al., 2023). The matrix contained 15,949,350 variants for 431 cultivars (Sarah et al., 2025). Missing values were imputed using Beagle software v.5.4 with default parameters (Browning et al., 2018). Genotyping data for 277 cultivars of the diversity panel present among the 431 cultivars were extracted from this matrix. PLINK software v2.0 (Chang et al., 2015) was used to retain only biallelic variants, both Single Nucleotide Polymorphisms (SNPs) and small insertions/deletions (indels), resulting in 14,153,069 positions, and to filter on minor allele frequency (MAF ≥ 5%), heterozygosity rate (Hz ≤ 0.6), Hardy-Weinberg Equilibrium (HWE test *P*-value ≥ 10^-20^), genotypic class frequency (GCf ≥ 0.02) and Linkage Disequilibrium (LD ≤ 0.99). We ended up with a matrix of 1,608,936 variants on 277 cultivars. Since only a subset of 241 and 246 cultivars were phenotyped in greenhouse and field environments, respectively, and to account for missing phenotyping values specific to each trait, we filtered again on MAF, before performing GWAS. The filtered matrix thus contained between 1,608,874 and 1,608,918 variants, depending on the trait and the environment considered.

### Detection of additive and non-additive QTLs with single- and multi-environment GWAS

In order to decipher the genetic determinism of carbon- and water-related traits in the grapevine diversity panel, GWAS analyses were performed following this strategy (Table 3):

**Table 3:** Summary of GWAS models performed in this study. “Traits” refers to BLUPs of the traits.

| Type of trait | Single- or multi-trait and environment | Method (R package) |
| --- | --- | --- |
| Carbon- and water-related traits | Single-trait<br>Single-environment | Additive and Non-additive effect using single-locus analyses (MM4LMM) + Local Score on single-locus results |
| Carbon- and water-related traits | Single-trait<br>Multi-environment | Additive effect single-locus analyses (metaGE and MTMM) + Local Score on single-locus results |
| Greenhouse PCA axes | Multi-trait from greenhouse environment<br>Single-environment | Additive and Non-additive effect using single-locus analyses (MM4LMM) + Local Score on single-locus results |
| Field PCA axes | Multi-trait from field environment<br>Single-environment | Additive and Non-additive effect using single-locus analyses (MM4LMM) + Local Score on single-locus results |
| Greenhouse + Field PCA axes | Multi-trait from both environments<br>Multi-environment | Additive and Non-additive effect using single-locus analyses (MM4LMM) + Local Score on single-locus results |

i/ Detection of additive and non-additive QTLs per environment (greenhouse or field) for each trait BLUP using a single-locus mixed-model implemented in R/MM4LMM v3.0.2 (Laporte et al., 2022). ii/ Detection of additive and non-additive QTLs using the coordinates from PCAs run either for each environment individually or on both environments, with a single-locus mixed-model (MM4LMM). iii/ Detection of multi-environment additive QTLs with two complementary approaches, the MTMM model (Korte et al., 2012) and a meta-analysis implemented in R/metaGE v1.1.0 (De Walsche et al., 2025), for the detection of QTLs with similar or opposite effects.

Finally, the *P*-values obtained from all single-locus mixed-models were used as input for a local score approach (Bonhomme et al., 2019). This method identified QTLs in which multiple linked genetic variants showed a consistent association with the phenotype, taking advantage of the high density of variants.

For more detail on GWAS models used and thresholds definition, see Supplementary Methods 3 and 4.

### QTL definition and gene annotation

In order to investigate candidate genes for the associated trait, we defined an interval around each significant or putative variant detected by the single-locus mixed-model approach or by the Local Score approach. This interval was based on the linkage disequilibrium (LD) decay estimated for each chromosome (for more detail on LD decay estimation, see Supplementary Method 5, Table S5). Overlapping intervals were merged between QTLs when LD between their most significant variants (topVariants) was > 0.8, with LD estimate being corrected by relatedness between cultivars and computed using R/LDcorSV v1.3.3 (Desrousseaux et al., 2020). After merging, all remaining distinct intervals were considered as QTL intervals.

We further investigated the genes overlapping these QTL intervals using the PN_T2T functional genome annotation v5.1 available in GRAPEDIA (https://grapedia.org/genomes/). We converted gene names from the PN_T2T (v5.1) genome to PN40024.v4 40X v4.3 (Velt et al., 2023) using the catalogue gene card from PlantaeViz (Navarro-Payá et al., 2022, https://plantaeviz.tomsbiolab.com/vitviz/catalogue/) and used the gene names from PN40024.v4 40X to assess their maximum expression in leaves, expressed in Reads Per Kilobase of transcript per Million reads mapped (RPKM), using GREAT (GRape Expression Atlas, which gathers published expression data) RNA-Seq data analysis workflow (https://great.colmar.inrae.fr/app/GREAT). Data were extracted with all filters set to “Select All” to keep all studies, except for the organ that was set to “Leaves”. To further identify the most likely candidate genes, we used the putative gene functions provided in the functional annotation v5.1 to screen the UniProt (https://www.uniprot.org/) and TAIR (https://www.arabidopsis.org/) databases for genes functions description.

## Acknowledgements

We thank the ETAP (LEPSE), M3P (LEPSE) and DAAV (AGAP Institut) team members for providing help during the preparation and phenotyping of the grapevine panel, especially Adrianus J. Westgeest, Jean-Pierre Péros, Timothée Flutre, Florent Pantin and Charles Romieu for their involvement in the early stages of the greenhouse experiment, and Maëlle Lis and Llorenç Cabrera-Bosquet for managing the PhenoArch platform. For genomic data, we thank Sylvain Santoni, Sandrine Dedet, Audrey Weber, Christine Tollon-Cordet, Valérie Laucou, Catherine Roux for sequence data generation and preprocessing. For the greenhouse experiment, we thank Freddy Gavanon and Mélyne Falcon for plant preparation and management. We thank Gaëlle Rolland, Thierry Pons and Raphaël Galtier for their help in leaf phenotyping. For the field experiment, we are grateful to the staff of INRAE Pech Rouge experimental vineyard, managed by Jean-Noël Lacapère, for maintenance of the diversity panel, and to Emmanuelle Garcia-Adrados for regular records of plant state; to the staff of Chambre d’Agriculture de l’Aude for grafting, plantation, and replacements; to Pierre François and Marc Farnos for helping with cutting harvest for grafting; to Thierry Lacombe for expert varietal identity checking. We are grateful to the staff of INRAE Vassal-Grape-BRC grapevine repository for long-term germplasm conservation and characterization, in particular to Cécile Marchal, Anne Mocoeur, Sandrine Dedet, Stéphanie Pattier and Anthony Chichrie. We thank Laurent Torregrosa for administration of ANR G^2^WAS project and Patrice This for administration of Défi clé Vinid’Occ.

## Author contributions

EC (collected data, curated data, managed field phenotyping, statistical analyses, writing original draft), SDE (collected data, preliminary analyses), RB (collected data, curated data), AD (designed and managed the greenhouse experiment, managed the diversity panel plantation in field, writing review and editing), TS (designed and managed the greenhouse experiment, writing review and editing), NS (managed the diversity panel growing in field), GS (sequence data preprocessing), BP (sequence data preprocessing, writing review), RB (project administration, writing review), DT (project administration, writing review), BP (supervised research, writing review and editing), VS (designed and managed phenotyping and statistical analyses, collected data, supervised research, writing review and editing) and AC-L (designed and managed phenotyping and analyses, collected data, supervised research, writing review and editing). All the authors have approved the manuscript.

## Data availability statement

The datasets supporting the conclusions of this article are available at the following private link in a public data repository (data.inrae.fr)(https://entrepot.recherche.data.gouv.fr/privateurl.xhtml?token=c913cbcc-8862-497d-87b9-c0decc030fe8) and will be made publicly available upon acceptance of the manuscript. The scripts for BLUP estimation and genomic analyses are publicly available in GitLab at https://forge.inrae.fr/eva.coindre/genomic_regions_leaf_traits_greenhouse_field_grapevine. Raw sequencing data can be accessed in ENA under the Project accession number PRJEB95058 and are described within the following datapaper https://hal.science/hal-05322783v1.

