## Supplementary_materials for "Genetic control of leaf carbon- and water-related traits in grapevine under greenhouse and field conditions"

**Supplementary Method 1: Leaf phenotyping protocol**

Transpiration rate (E, mmol $m^{-2}$ $s^{-1}$), stomatal conductance ($g_s$, mol $m^{-2}$ $s^{-1}$), quantum efficiency of photosystem II in light (ΦPSII) and electron transport rate (ETR, µmol $m^{-2}$ $s^{-1}$) were phenotyped using the Li600 hand-held poro-fluorometer. Two measurements were taken on each side of the principal vein on the abaxial surface of each leaf measured, and were averaged. A proxy of chlorophyll content was assessed as SPAD value using SPAD-502 Chlorophyll Meter (Konica-Minolta, Tokyo, Japan). Measurements were taken on four different spots of each leaf measured, avoiding principal veins, and averaged to obtain a single value for each leaf. A micro-portable near infrared spectrometer, the MicroNIR (MicroNIR™ Onsite-W, VIAVI, Chandler, AZ, United States) was also used *in situ* to collect two spectra per leaf (one for each side of the leaf) in the spectral range from 908 nm to 1,676 nm with 125 wavelengths. The two spectra were averaged to obtain a single spectrum for each leaf.

For a subset of 102 leaves and 37 leaves (corresponding to 31 and 37 different cultivars randomly selected, respectively) from the greenhouse and the field experiment, respectively, net $CO_2$ assimilation rate ($A_n$, µmol $m^{-2}$ $s^{-1}$) was also measured (Table S1) using an InfraRed Gas-exchange Analyzer (IRGA) equipped with a light-emitting diode (Li6800, Portable Photosynthesis System, Licor, Lincoln, United States). The $CO_2$ condition in the leaf chamber was set to 400 µmol $mol^{-1}$ and for each measured leaf, the actinic light was adjusted to a value corresponding to the PPFD measured simultaneously by the Li600 device.

Following these *in situ* measurements on leaves, a 3 cm diameter disk was sampled for each of the 1,449 greenhouse leaves and 1,102 field leaves. These disks were immediately weighed after sampling for all greenhouse leaves and for the subset of 37 field leaves to obtain their fresh weight (FW, mg) with 0.1 mg accuracy (Sartorius balance, QUINTIX224-1S, Göttingen, Germany). For both experiments, all disks sampled were oven-dried for one week at 60°C and weighed again to obtain their dry weight (DW, mg). Leaf Mass per Area (LMA, mg $cm^{-2}$) was then calculated as $\frac{DW}{disk\ area}$ (with disk area = 7.07 $cm^2$), Water Quantity (WQ, mg $H_2O$ per leaf disk) as $FW - DW$ and Water Content ($WC_f$, mg $H_2O$ $mg^{-1}$ fresh weight) as $\frac{WQ}{FW}$. Finally, all 1,449 and 1,102 dried disks from greenhouse and field experiments, respectively, were used to collect four additional near infrared spectra per disk, which were then averaged, using a laboratory spectrometer ASD (ASD LabSpec® 4, Analytical Spectral Devices, Boulder, USA) that covers a spectral range from 350 nm to 2,500 nm with 2,150 wavelengths.

**Supplementary Method 2: Trait prediction with PLSR models**

$A_n$ was predicted using the measurements of the poro-fluorometer Li600 device. For the 1410 greenhouse leaves, the PLSR model had previously been built and validated on the 102 leaves where $A_n$ was measured with the Li6800 device (Table S1) (Coindre et al., 2026). For the field leaves, a PLSR model was built combining measures from both Li600 and Li6800 devices on 157 leaves of other field environments conducted by the team, using the exact same phenotyping process. This model was validated on 37 leaves from the 2024 field experiment (Table S2) and applied on the 1102 leaves to be predicted (Table S1).

LMA, $WC_f$ and WQ were not predicted in the greenhouse because they were already measured on all 1449 leaves (Table S1). For the 1102 leaves from the field, they were predicted using a PLSR model built with 1410 measures from the 2022 greenhouse experiment and 232 leaves from other potted plants in outdoor environments (Table S2) conducted by the team and using the exact same phenotyping process. We increased the number of calibration samples with leaf disks from outdoor environments to ensure that they were representative of the climatic conditions of the field samples to be predicted. The PLSR model combined these measurements with Near InfraRed Spectroscopy (NIRS) high-throughput data. A 4-fold cross-validation was performed to determine the optimal number of latent variables (LVs), as the number of LVs that minimized the average root mean squared error (RMSE), with a maximum set to 20. The resulting model was validated on 37 leaves from the field experiment on which LMA, $WC_f$ and WQ were measured (Table S2) before being applied for the prediction of the 1102 leaves from the field experiment (Table S1).

PLSR models were considered acceptable when the coefficient of determination of the validation set ($R^2_{val}$) exceeded 0.5 and its normalized root mean squared error ($NRMSE_{val} = \frac{RMSE_{val}}{\bar{x}}$ with $\bar{x}$ the mean of the trait) was below 15%.

**Supplementary Method 3: GWAS models**

Additive QTLs were identified using a single-locus mixed-model, based on (Yu et al., 2006) and implemented in R/MM4LMM:

$$\mathbf{y} = \mu + \mathbf{x}\beta + \mathbf{u} + \boldsymbol{\varepsilon} \quad \text{(eq 2)}$$

Where **y** was the vector of BLUPs, μ was the overall mean, **x** was the vector of genotypic data at the variant studied (coded in allelic doses of the reference allele, 0/1/2), β was the corresponding additive effect, **u** was the vector of random polygenic effects, which followed a normal distribution $N(0, K\sigma_u^2)$ with K the covariance matrix and $\sigma_u^2$ the genetic variance, and **ε** was the vector of random residual effects, which followed a normal distribution $N(0, I\sigma_\varepsilon^2)$ with I the identity matrix and $\sigma_\varepsilon^2$ the residual variance. The covariance matrix K (kinship) was calculated with variant weights based on local linkage disequilibrium, as implemented in LDAK software v5.2 (Speed et al., 2012). The genomic heritability was calculated as $h_{genomic}^2 = \frac{\sigma_u^2}{\sigma_u^2+\sigma_e^2}$ with $\sigma_u^2$ and $\sigma_e^2$ extracted from a reduced version of equation 2, without the fixed variant term:

$$\mathbf{y} = \mu + \mathbf{u} + \boldsymbol{\varepsilon} \text{ (eq 3)}$$

In order to specifically detect non-additive QTLs, the single-locus mixed-model (equation 2) was adapted by converting variant information into factors and then performing a test in an analysis of variance (ANOVA) manner.

In order to identify similar and independent associations between both environments, a single-locus multi-environment GWAS was performed by using the multi-trait mixed-model (MTMM) approach (mtmm.gei.gwas.r function available at https://forgemia.inra.fr/vincent.segura/sybiopop_chi.git):

$$y = \begin{bmatrix} y_G \\ y_F \end{bmatrix} = s_G\mu_G + s_F\mu_F + \mathbf{x}\beta + (\mathbf{x} \times \boldsymbol{s_G})\alpha + \mathbf{u} + \boldsymbol{\varepsilon} \text{ (eq 4)}$$

Where $\boldsymbol{y_G}$ and $\boldsymbol{y_F}$ were the vectors of BLUPs from greenhouse and field environments respectively, $\boldsymbol{s_i}$ was a vector containing 1 for all values belonging to environment i (with i = G for greenhouse and i = F for field) and 0 otherwise, $\mu_G$ and $\mu_F$ were the overall means of greenhouse and field environments, respectively, **x** and β were the same as in equation 2 (except for the dimension of the vector **x**), α was the vector of interaction effects between variants and environments, **u** was a vector of random polygenic effects which followed a normal distribution N(0, $\sum_C \otimes$K), with $\sum_C$ the genetic variance-covariance matrix between both environments and K the kinship matrix, **ε** was the vector of random residual effects which followed a normal distribution N(0, $\sum_\varepsilon \otimes$I), with $\sum_\varepsilon$ the residual variance-covariance matrix between both environments and I the identity matrix. Since we do not expect any residual covariance between environments, the residual covariance between environments was set to 0. The genetic and residual variance-covariance matrices were first estimated in a null model (without any marker effect) using R/breedR v0.12-5 (Muñoz & Sanchez, 2020). Then variant effects were estimated using the Generalized Least Squares approach, identifying (i) both similar and interaction effects between environments (“full model”), (ii) only similar effects between environments (“common model”) and (iii) effects that vary between environments (“gei model”).

In complement of the MTMM model, which explicitly models the covariance between environments, meta-analyses combining the results of environment-specific GWAS (using *P*-values and effects of the single-locus mixed-models for each environment), were also performed to detect consistent effects across environments, independent of environment-specific variance, as implemented in R/metaGE (De Walsche & Mary-Huard, 2025) (Table 3). We used the fixed procedure that considered the QTL to have the same effect across environments to identify similar QTLs across environments. The model can be described as follows:

$$Z_m = X\mu_m + E_m \quad \text{(eq 5)}$$

where $Z_m$ was the vector of Z-scores for marker m over a set of *K* different GWASs performed in individual environments, X was an incidence matrix of size $K \times J$ that contains 1 if environment k belongs to group j and 0 otherwise, $\mu_m$ was the group-specific marker effects's vector. $E_m$ was the vector containing the residual error, which followed a normal distribution N($0_K$, $\sum_m$) with $\sum_m$ the $K \times K$ inter-environment correlation matrix. In the case of a stable allelic effect across environments (i.e. $J = 1$), the model tested was reduced to $H_0$:$\{\mu_m = 0\}$.

For all GWAS models, a variant was considered as significantly associated with the genotypic values when its *P*-value of association was lower than the Bonferroni threshold calculated as ($\frac{0.05}{nb\ of\ variants} \approx 3.1 \times 10^{-8}$, corresponding to a -$\log_{10}$(*P*-value)≈7.5). A variant was considered as putatively associated when its *P*-value was lower than $10^{-6}$.

Finally, the Local Score approach was applied to the *P*-values from all single-locus mixed-models. The tuning parameter $\xi$ was fixed at 3, so only variant with a -$\log_{10}$(*P*-value) larger than 3 were considered positively in the score calculation and cumulated using the Lindley process (Bonhomme et al., 2019; Fariello et al., 2017). The threshold used to identify putative variants was fixed at a Lindley score of 9.

The percentage of variance explained (PVE) was calculated for each topVariant in QTLs detected by single-trait and single-environment methods as follows (Nagelkerke, 1991):

$$PVE = 1 - exp\left(-\frac{2}{n}(l_1 - l_0)\right) \quad \text{(eq 6)}$$

where $l_1$ was the log-likelihood of the mixed model including the topVariant, $l_0$ was the log-likelihood of the null model, without the topVariant effect and n was the number of cultivars. PVE cannot be estimated for multi-environment topVariants.

**Supplementary Method 4: Thresholds definition for GWAS analyses**

For the single-locus GWAS model, a variant was considered significantly associated with the trait when the *P*-value from the Wald test statistic was smaller than a defined threshold controlling the family-wise error-rate at 5% to account for multiple testing. The high coverage density of variants in the resequencing matrix, with a marker every 310 bp approximately, resulted in a Bonferroni threshold of -$\log_{10}$($\frac{0.05}{nb\ of\ variants}$) ≈ 7.5. Previous grapevine studies calculated a Bonferroni threshold on a subset of nearly independent Single-Nucleotide polymorphisms (SNPs) based on a linkage disequilibrium (LD) pruning method (Coupel-Ledru et al., 2024; Chir et al., 2025) resulting in a threshold close to 5.5. In this study, the high density of markers did not allow for the calculation of a nearly independent variant subset because of the complexity of parameter choice. In order to avoid missing true positive QTLs, we decided to set a second threshold with *P*-value lower than $10^{-6}$ ($-\log_{10}(P\text{-value})=6$), higher than those calculated in (Coupel-Ledru et al., 2024; Chir et al., 2025).

The Local Score approach was initially developed to increase the detection power of selection signals through genome scans (Fariello et al., 2017). This approach was then translated to improve the detection of QTLs with small effects by using *P*-values from GWAS models (Bonhomme et al., 2019). Here the tuning parameter $\xi$ was set to 3, which corresponds to a *P*-value threshold of $10^{-3}$ for single-locus tests, and computed significant thresholds based on a re-sampling approach of score sequences *M* of fixed length (from 10,000 to 70,000 bp), coupled with autocorrelation $\rho$ (from 0.025 to 0.275). Each pair *M* x $\rho$ was repeated 5,000 times to estimate the coefficients of the Gumbel distribution (Fariello et al., 2017). Threshold estimates, with a type I error rate of 5%, were computed for each of the 19 chromosomes for 23 randomly selected traits. To limit the discovery of false QTLs and enable comparison between traits, we decided to set the threshold using the 95$^{th}$ percentile of the distribution of all thresholds over each trait and each chromosome, which resulted in a Lindley score of 9.

**Supplementary Method 5: Linkage disequilibrium decay estimation**

The interval around each significant or putative variant was determined using the LD decay extent estimated for each chromosome. For that, 100,000 pairs of randomly selected variants were sampled from each chromosome. LD was estimated by calculating the $r^2_V$ estimator (Mangin et al., 2012), the $r^2$ corrected by relatedness (kinship) between cultivars, as implemented in R/LDcorSV v1.3.3 (Desrousseaux et al., 2020). The kinship matrix was estimated as a modified kinship matrix with variants weighted according to local LD, using LDAK software v5.2 (Speed et al., 2012). For each chromosome, LD maximum value (Y-intercept) and decay were estimated using the mathematical function proposed by Hill and Weir (Hill & Weir, 1988) and LD decay extent was defined as the distance at which LD had decreased to half its maximum (Table S5).

**Table S1: Number of measured and predicted samples for each trait according to the environment.**

LMA: Leaf mass per area, $WC_f$: Water content, WQ: Water quantity, SPAD: Proxy of chlorophyll content, $g_s$: Stomatal conductance, E: Transpiration rate, ETR: Electron transport rate, ΦPSII: Quantum efficiency of photosystem II in light, $A_n$: Net $CO_2$ assimilation rate and $WUE_{intr\_cal}$: Intrinsic water use efficiency. $WUE_{intr\_cal}$ was calculated by dividing the predicted values of $A_n$ by the measured values of $g_s$. In the greenhouse environment, $WUE_{intr\_cal}$ was calculated only for plants under well-watered and moderate water deficit conditions.

| **Trait** | **Greenhouse** | | | **Field** | | |
|---|---|---|---|---|---|---|
| | **Measured** | **Predicted** | **Calculated** | **Measured** | **Predicted** | **Calculated** |
| LMA | 1449 | - | - | 37 | 1102 | - |
| $WC_f$ | 1449 | - | - | 37 | 1102 | - |
| WQ | 1449 | - | - | 37 | 1102 | - |
| SPAD | 1444 | - | - | 1102 | - | - |
| ETR | 1445 | - | - | 1102 | - | - |
| ΦPSII | 1446 | - | - | 1102 | - | - |
| $g_s$ | 1446 | - | - | 1102 | - | - |
| E | 1446 | - | - | 1102 | - | - |
| $A_n$ | 102 | 1410 | - | 37 | 1102 | - |
| $WUE_{intr\_cal}$ | 102 | - | 913 | 37 | - | 1102 |

**Table S2: Number of leaves used to calibrate and validate PLSR model for each trait with the corresponding device, for prediction on samples of the 2024 field environment.**

$A_n$: Net $CO_2$ assimilation rate, LMA: Leaf mass per area, $WC_f$: Water content, WQ: Water quantity.

| | | **Calibration samples** | | | **Validation samples** |
|---|---|---|---|---|---|
| **Trait** | **Device** | **Greenhouse environment 2022** | **Other outdoor potted environment** | **Other field environments** | **Field environment (2024)** |
| $A_n$ | Li600 | - | - | 157 | 37 |
| LMA | ASD | 1410 | 232 | - | 37 |
| $WC_f$ | MicroNIR | 1410 | 232 | - | 37 |
| WQ | MicroNIR | 1410 | 232 | - | 37 |

**Table S3: PLSR features obtained for the validation set (*n*=37) for each predicted trait in the field environment.**

$A_n$: Net $CO_2$ assimilation rate, LMA: Leaf mass per area, $WC_f$: Water content, WQ: Water quantity. $R^2_{val}$: coefficient of determination of the validation set. $RMSE_{val}$: root mean squared error of the prediction (in the trait unit). $NRMSE_{val}$: normalized root mean squared error of the prediction (in percentage). $Bias_{val}$: average difference between predicted and measured values. The device in bold corresponds to the one retained for the prediction of the corresponding trait for all field leaves.

| **Trait** | **Device** | $\mathbf{R^2_{val}}$ | $\mathbf{RMSE_{val}}$ | $\mathbf{NRMSE_{val}}$ | $\mathbf{Bias_{val}}$ |
|---|---|---|---|---|---|
| $A_n$ | **Li600** | 0.67 | 1.7 | 11% | -0.06 |
| LMA | MicroNIR | 0.81 | 0.48 | 7.9% | 0.27 |
| | **ASD** | 0.87 | 0.42 | 6.9% | -0.19 |
| $WC_f$ | **MicroNIR** | 0.83 | 0.01 | 1.4% | 0.01 |
| | ASD | 0.29 | 0.04 | 5.6% | 0.04 |
| WQ | **MicroNIR** | 0.56 | 9.5 | 9.4% | -5.29 |
| | ASD | 0.18 | 23.1 | 22.9% | 20.78 |

**Table S4: Significant fixed effects retained in the linear mixed-model for the extraction of genotypic BLUPs for each trait in greenhouse and field environments.**

LMA: Leaf mass per area, $WC_f$: Water content, WQ: Water quantity, SPAD: Proxy of chlorophyll content, $g_s$: Stomatal conductance, E: Transpiration rate, ETR: Electron transport rate, ΦPSII: Quantum efficiency of photosystem II in light, $A_n$: Net $CO_2$ assimilation rate and $WUE_{intr\_cal}$: Intrinsic water use efficiency. Square-root transformation was applied to WQ, $g_s$ and E and log transformation (base 10) was applied to $WUE_{intr\_cal}$ before adjusting the linear mixed-model, as the data were not normally distributed. Date is the date of measurement and time the time of measurement. $T_{air}$ is the mean air temperature. PPFD is the mean Ambient Photosynthetically Photon Flux Density. VPD is the mean vapor pressure deficit. $T_{air}$, PPFD and VPD were recorded with probes in the greenhouse environment and with the Li600 device in the field environment. Water treatment is the water treatment applied to the plant in the greenhouse environment. Exp. duration is the duration in days between the entrance of the plant into the greenhouse and measurement. Density is the plant density around the plant in the greenhouse environment. Last irrigation is the time elapsed between the last irrigation of the plant and measurement in the greenhouse environment. Block corresponds to the randomized complete block in the field environment, row and position in the row is the position of the unit plot in the corresponding row in the field environment. Cells are filled in gray if the fixed effect is not relevant to the corresponding environment. X corresponds to fixed effects retained in the linear mixed-model.

| Trait | Unit | Transformation applied to data | Environment | Date | Time | $T_{air}$ | PPFD | VPD | Water treatment | Exp. duration | Density | Last irrigation | Block | Row | Position in the row |
|---|---|---|---|---|---|---|---|---|---|---|---|---|---|---|---|
| LMA | mg $cm^{-2}$ | none | Greenhouse | X | X | | | | X | X | | | | | |
| | | | Field | X | | | | | | | | | X | X | X |
| $WC_f$ | mg $H_2O$ $mg^{-1}$ fresh weight | none | Greenhouse | X | X | X | | | X | | X | | | | |
| | | | Field | X | X | | | | | | | | X | X | X |
| WQ | mg $H_2O$ | Square-root | Greenhouse | X | | | X | | X | | | | | | |
| | | | Field | X | X | | | | | | | | X | X | X |
| SPAD | SPAD unit | none | Greenhouse | X | | | | | X | X | | | | | |
| | | | Field | X | | | | | | | | | X | X | X |

| Trait | Unit | Transformation applied to data | Environment | Date | Time | $T_{air}$ | PPFD | VPD | Water treatment | Exp. duration | Density | Last irrigation | Block | Row | Position in the row |
|---|---|---|---|---|---|---|---|---|---|---|---|---|---|---|---|
| ETR | µmol $m^{-2}$ $s^{-1}$ | none | Greenhouse | X | X | | | | X | | | X | | | |
| | | | Field | X | | X | X | | | | | | X | X | |
| ΦPSII | NA | none | Greenhouse | X | X | | | | X | | | X | | | |
| | | | Field | X | | X | X | | | | | | X | X | |
| $g_s$ | mol $m^{-2}$ $s^{-1}$ | Square-root | Greenhouse | X | | | | | X | | | | | | |
| | | | Field | X | | X | X | X | | | | | X | X | |
| E | mmol $m^{-2}$ $s^{-1}$ | Square-root | Greenhouse | X | X | | | | X | | | | | | |
| | | | Field | X | | X | X | X | | | | | X | X | |
| $A_n$ | µmol $m^{-2}$ $s^{-1}$ | none | Greenhouse | X | X | | | | X | | | X | | | |
| | | | Field | X | | X | X | X | | | | | X | X | |
| $WUE_{intr\_cal}$ | µmol $CO_2$ $mol^{-1}$ $H_2O$ | $Log_{10}$ | Greenhouse | X | | | | | X | | | | | | |
| | | | Field | X | | X | X | X | | | | | X | X | |

**Table S5: Extent of the linkage disequilibrium (LD) decay for each chromosome in base pairs (bp).**

| Chromosome | LD decay extent (in bp) |
|---|---|
| Calculated over whole genome | 63,998 |
| 1 | 56,474 |
| 2 | 186,182 |
| 3 | 39,121 |
| 4 | 133,767 |
| 5 | 43,026 |
| 6 | 56,009 |
| 7 | 44,750 |
| 8 | 90,645 |
| 9 | 44,235 |
| 10 | 46,304 |
| 11 | 62,190 |
| 12 | 34,983 |
| 13 | 36,860 |
| 14 | 64,844 |
| 15 | 62,053 |
| 16 | 24,827 |
| 17 | 148,129 |
| 18 | 55,006 |
| 19 | 55,094 |

**Table S6: List of the QTLs found for each trait, according to the environment (greenhouse or field), the GWAS method (MM4LMM, metaGE, MTMM and Local Score) and the variant effect modeling used (additive or ANOVA).**

This supplementary table is available at the following link: https://entrepot.recherche.data.gouv.fr/privateurl.xhtml?token=c913cbcc-8862-497d-87b9-c0decc030fe8

The lines written in red correspond to the subset of 10 QTLs presented in Table 2.

Each block of lines (shaded or unshaded) corresponds to all significant or putative variants detected in an environment for a specific trait, method and model. These variants have overlapping intervals and are therefore grouped under the same QTL. The first column 'QTL.id' indicates the unique name of the QTL. 'Trait' indicates the associated trait. 'Environment' refers to the environment in which the QTL was found, either in greenhouse or in field, or in both in multi-environment analyses. 'Method' corresponds to the GWAS method used to detect the QTL, either with $-\log_{10}(P) > 6$ for MM4LMM, metaGE and MTMM single-locus association tests, or with a Lindley score > 9 using the Local Score approach based on the *P*-values of MM4LMM, metaGE, and MTMM. 'Model' corresponds either to additive (add) or ANOVA variant effect modeling (anova). 'TopVariant' is the name (chromosome:position) of the most significant variant in the QTL for the given trait. 'Chr' indicates chromosome number. 'Interval start' and 'Interval end' correspond to the start and end of the interval defined around the topVariant for the given trait. '-log10(P) add (MM4LMM)' and '-log10(P) anova (MM4LMM)' are the -log10(P-value) for the MM4LMM single-locus mixed-model at the topVariant with the additive and ANOVA variant models, respectively. '-log10(P) (metaGE)', '-log10(P) (MTMM full)', '-log10(P) (MTMM common)' and '-log10(P) (MTMM gei)' are the -log10(P-value) for the additive model of metaGE, MTMM full, MTMM common and MTMM gei multi-environment single-locus mixed-model at the topVariant, respectively. 'Lindley score add' and 'Lindley score anova' are the maximum Lindley score values for the intervals of the QTLs detected by the local score approach using the single-locus mixed-model (MM4LMM, metaGE, MTMM) with additive or ANOVA model, respectively. 'PVE' is the proportion of BLUP variance explained by the topVariant of the QTL, expressed in percentage and calculated for single-environment topVariant only. 'MAF' is the minor allele frequency of the topVariant.

**Table S7: List of genes underlying the main QTLs of interest.**

This supplementary table is available at the following link: https://entrepot.recherche.data.gouv.fr/privateurl.xhtml?token=c913cbcc-8862-497d-87b9-c0decc030fe8

Genes were searched for the 10 QTLs of particular interest presented in Table 2, within an interval defined around the variants detected by the single-locus mixed-model approach or by the Local Score approach, based on the linkage disequilibrium (LD) decay extent estimated for each chromosome. Best candidate genes are highlighted in red. The first column 'QTL.id' indicates the unique name of the QTL.

'vitvi.gene.id_v5' is the gene name based on the *Vitis vinifera* reference genome PN_T2T (v5) (Shi et al., 2023). 'vitvi.gene.id_v4.3' is the gene name based on the *Vitis vinifera* reference genome PN40024.v4 40X (v4.3) (Velt et al., 2023). 'Chr' indicates the chromosome number. 'start_T2T.v5' and 'end_T2T.v5' are the gene start and end position based on the *Vitis vinifera* reference genome PN_T2T (v5) (Shi et al., 2023). 'refseq.genedesc' is the putative function of the gene retrieved from the PN_T2T annotation (v5.1) (Shi et al., 2023), when available. 'full.description' is the complete description of the gene function, retrieved from TAIR and UniProt, when available. 'max.expression.leaves' is the maximum expression of the gene in the leaves, expressed in Reads Per Kilobase of transcript per Million reads mapped (RPKM), retrieved from GREAT, from all RNA-seq data available on the platform for leaves samples.

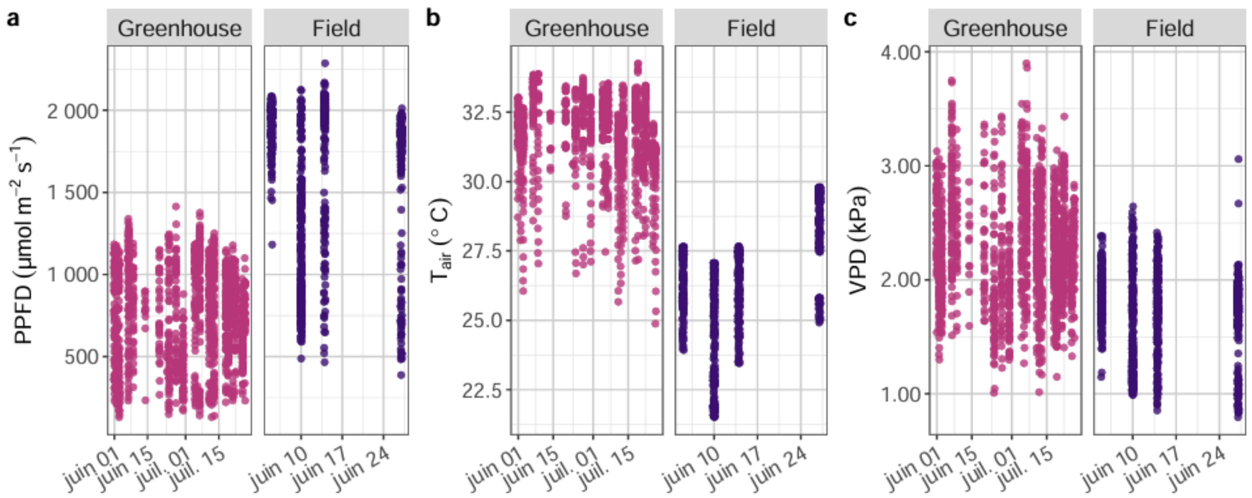


**Figure S1: Distribution of climatic conditions during leaf traits measurements in the greenhouse and the field environments.**

**a** Photosynthetically photon flux density (PPFD), **b** Air temperature ($T_{air}$) and **c** Leaf vapor pressure deficit (VPD).

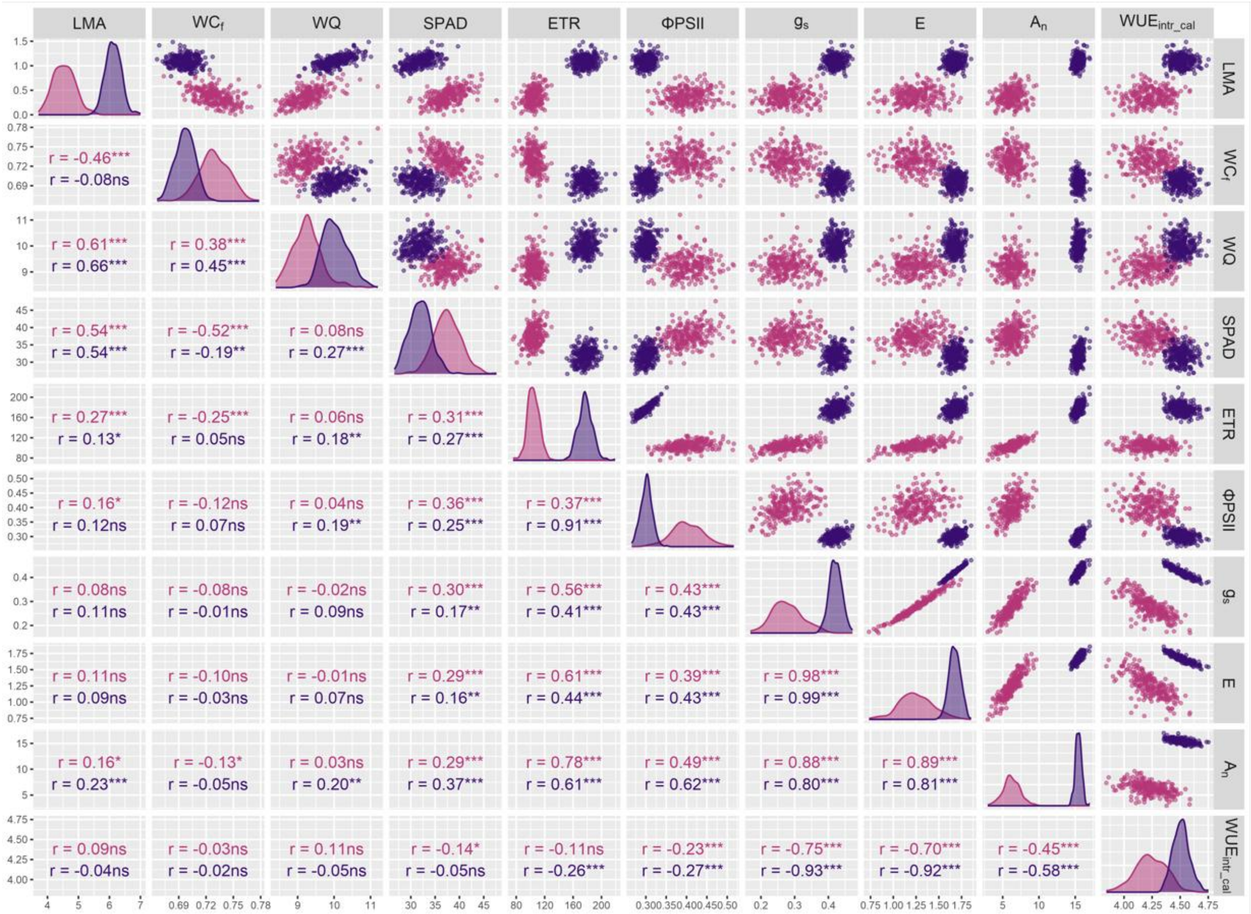


**Figure S2: Genetic correlation coefficients and scatter plots between BLUP traits, colored by environment.**

The diagonal shows the distribution of the trait genotypic BLUPs according to the environment. The upper part shows scatter plots between pairs of traits. The lower part gives Pearson correlation coefficients between pairs of traits with significance level. Pink is for the greenhouse environment and purple for the field environment.

Significance level is indicated for each correlation as follows: * = *P*-value ≤ 0.05; ** = *P*-value ≤ 0.01; *** = *P*-value ≤ 0.001. LMA: Leaf mass per area (in mg $cm^{-2}$), SPAD: Proxy of chlorophyll content, $WC_f$: Water content (in mg $H_2O$ $mg^{-1}$ fresh weight), WQ: Water quantity (in mg $H_2O$ per leaf disk), ETR: Electron transport rate (in µmol $m^{-2}$ $s^{-1}$), ΦPSII: Quantum efficiency of photosystem II in light, $g_s$: Stomatal conductance (in mol $m^{-2}$ $s^{-1}$), E: Transpiration rate (in mmol $m^{-2}$ $s^{-1}$), $A_n$: Net $CO_2$ assimilation rate (in µmol $m^{-2}$ $s^{-1}$) and $WUE_{intr\_cal}$: Intrinsic water use efficiency (in µmol $CO_2$ $mol^{-1}$ $H_2O$). Square-root transformation was applied to WQ, $g_s$ and E and log transformation (base 10) was applied to $WUE_{intr\_cal}$ before adjusting the linear mixed-model, as the data were not normally distributed.

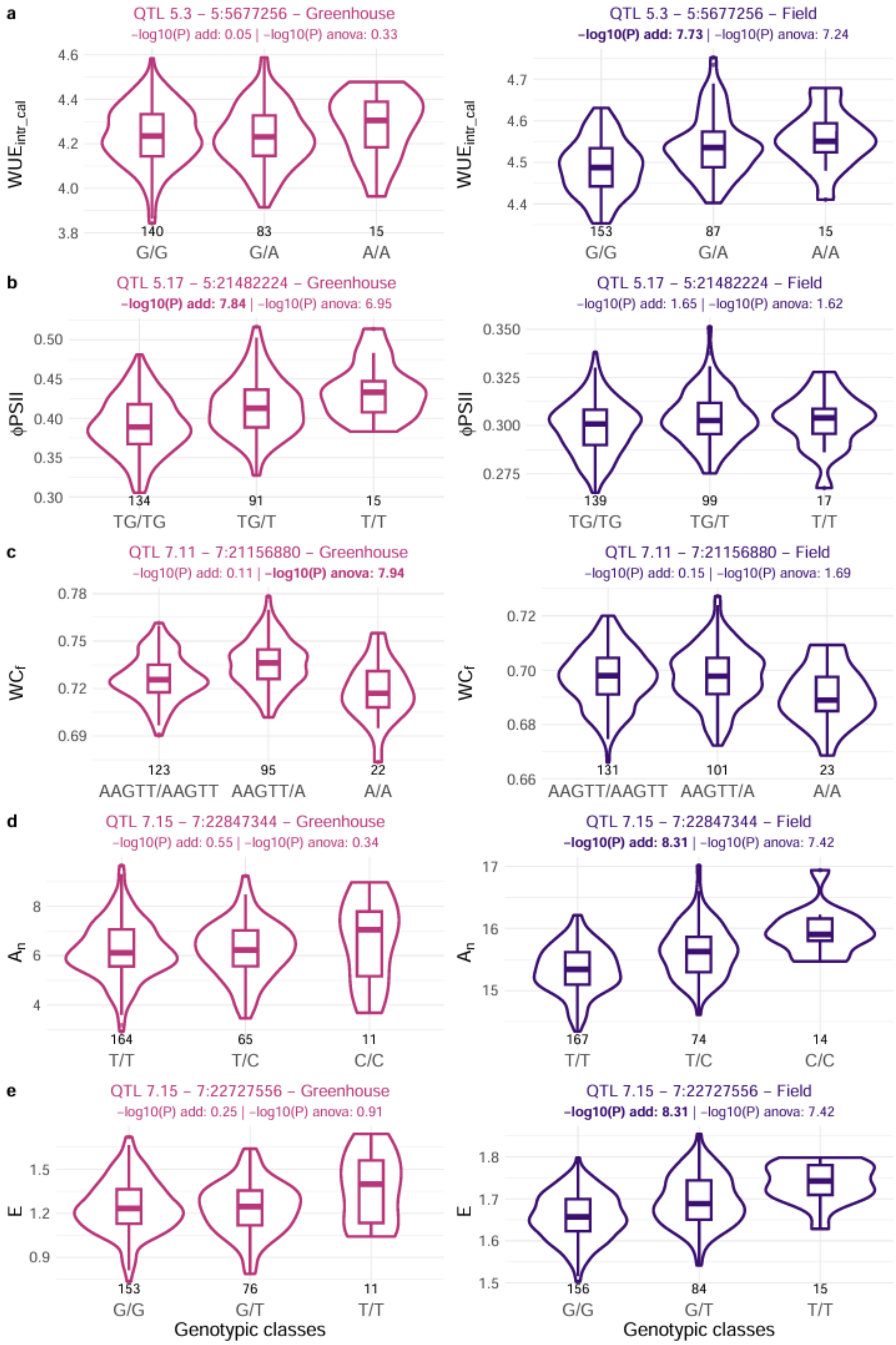


**Figure S3: Boxplots and violinplots showing the effect of genotypic classes on trait BLUPs in greenhouse and field environments, for topVariants passing the Bonferroni threshold in only one environment with the single-environment single-locus mixed-model (Criteria 1, Table 2).**

These results are shown in Figure 4 for QTL 7.2 for electron transport rate (ETR).

**a** is the topSNP of the QTL 5.3 for intrinsic water use efficiency ($WUE_{intr\_cal}$ in µmol $CO_2$ $mol^{-1}$ $H_2O$), **b** is the topVariant (small indel) of the QTL 5.17 for quantum efficiency of photosystem II in light (ΦPSII), **c** is the topVariant (small indel) of the QTL 7.11 for water content ($WC_f$ in mg $H_2O$ $mg^{-1}$ fresh weight), **d** and **e** are the topSNPs of the QTL 7.15 for net $CO_2$ assimilation rate ($A_n$ in µmol $m^{-2}$ $s^{-1}$) and transpiration rate (E in mmol $m^{-2}$ $s^{-1}$), respectively. The number of cultivars in each genotypic class is given under the corresponding boxplot. -$log_{10}$(P) add is the -$log_{10}$(*P*-value) for the additive only model and -$log_{10}$(P) anova is the -$log_{10}$(*P*-value) for the additive + non-additive model. -$log_{10}$(*P*-value) > $-log_{10}(\frac{0.05}{nb\ of\ variants}) = 7.5$ are highlighted in bold. Square-root transformation was applied to E and log transformation (base 10) was applied to $WUE_{intr\_cal}$ before adjusting the linear mixed-model, as the data were not normally distributed.

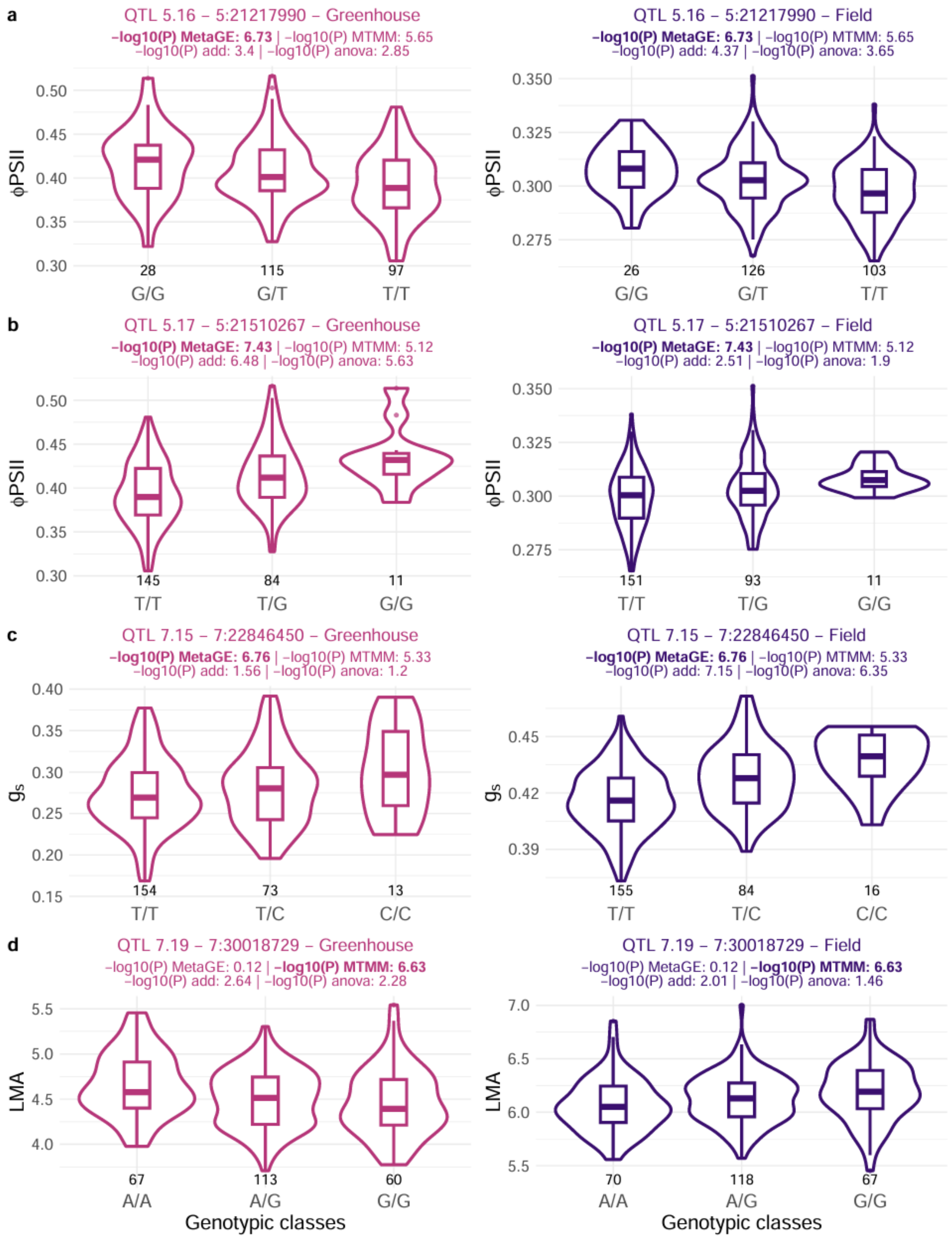


**Figure S4: Boxplots and violinplots showing the effect of genotypic classes on trait BLUPs (in each greenhouse and field environment), for each topSNP of QTLs detected by multi-environment mixed-model (Criteria 2 or 3, Table 2).**

These results are shown in Figure 5 for QTL 11.4 for net $CO_2$ assimilation rate and Figure 6 for QTL 16.4 for net $CO_2$ assimilation rate.

**a** TopSNP of QTL 5.16 for quantum efficiency of photosystem II in light (ΦPSII), **b** TopSNP of QTL 5.17 for quantum efficiency of photosystem II in light (ΦPSII), **c** TopSNP of QTL 7.15 for transpiration rate (E in mmol $m^{-2}$ $s^{-1}$) and **d** TopSNP of QTL 7.19 for leaf mass per area (LMA in mg $cm^{-2}$). The number of cultivars in each genotypic class is given under the corresponding boxplot. $-log_{10}$(P) MetaGE is the $-log_{10}$(*P*-value) for the meta-analysis model and $-log_{10}$(P) MTMM is the $-log_{10}$(*P*-value) for the MTMM approach. $-log_{10}$(P) add is the $-log_{10}$(*P*-value) for the additive only model and $-log_{10}$(P) anova is the $-log_{10}$(*P*-value) for the additive + non-additive model using the single-environment single-locus mixed-model. *P*-value$<10^{-6}$ are highlighted in bold. Square-root transformation was applied to E before adjusting the linear mixed-model, as the data were not normally distributed.

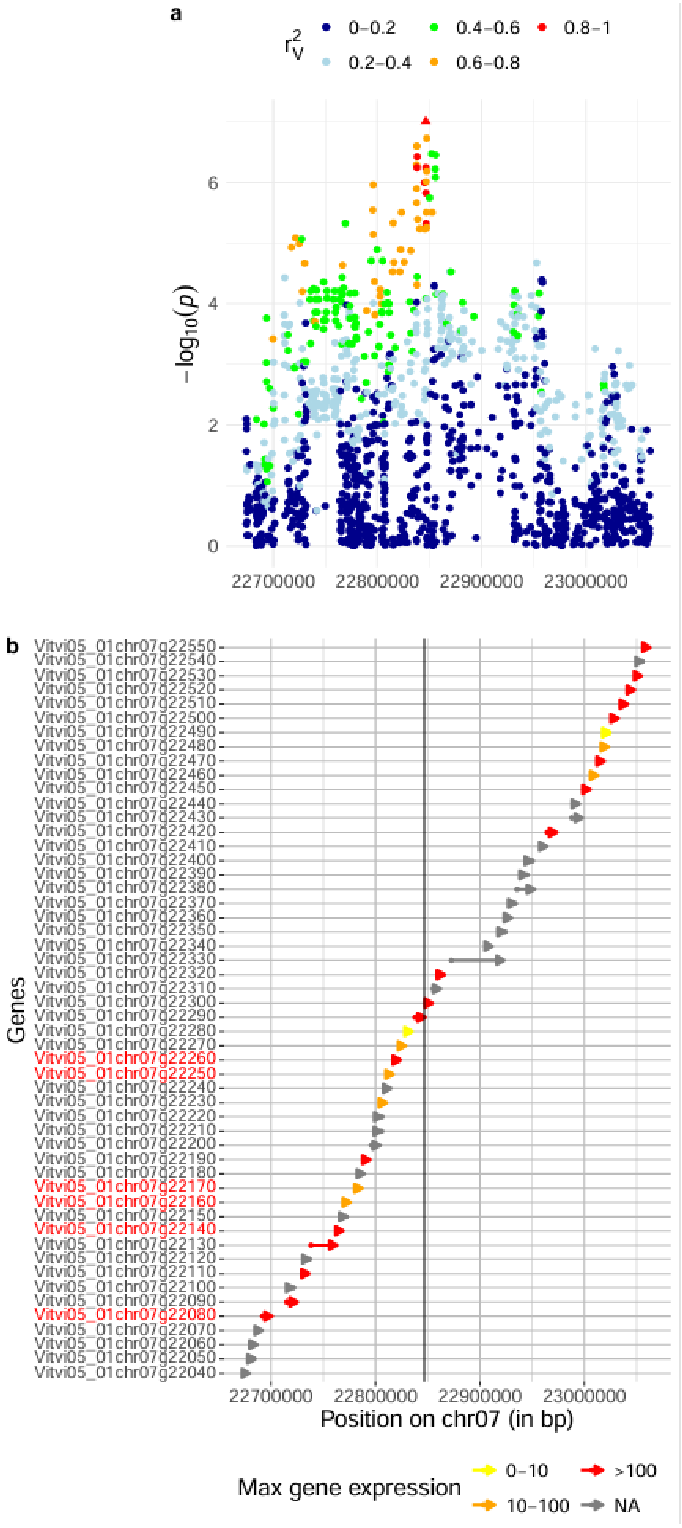


**Figure S5: Fine characterisation of QTL 7.15 which presented a similar effect between both greenhouse and field environments for genotypic values (BLUPs) of net $CO_2$ assimilation rate ($A_n$), transpiration rate (E), stomatal conductance ($g_s$) and intrinsic water use efficiency ($WUE_{intr\_cal}$).**

**a** Zoom of a genome-wide Manhattan plot performed using a meta analysis using transpiration rate from both greenhouse and field environments. Colors correspond to the local linkage disequilibrium corrected by the relatedness of cultivars ($r^2_V$) based on the topSNP 7:22846450 (triangle). **b** Maximum gene expression in leaves (color of the arrows) extracted from GREAT (GRape Expression Atlas) RNA-Seq data analysis workflow for the “leaf” organ across all studies, expressed in Reads Per Kilobase of transcript per Million reads mapped (RPKM). The black line corresponds to the position of the topSNP, aligned with **a**. Genes with red names are those with a putative function related to leaf polarity development and stomatal control. The size of the arrow is proportional to the size of the gene.

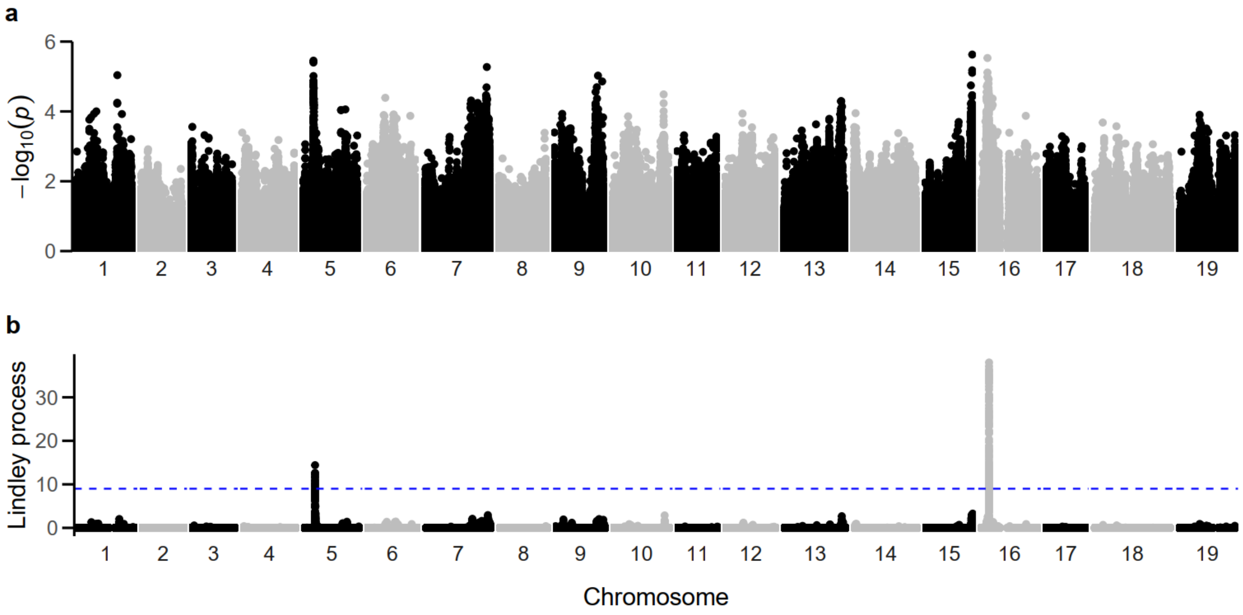


**Figure S6: a Manhattan and b local score plots for the gei model of the MTMM method performed for genotypic values (BLUPs) of net $CO_2$ assimilation rate ($A_n$) from greenhouse and field environments.**

The blue dashed line is the Local Score threshold (Lindley score of 9).

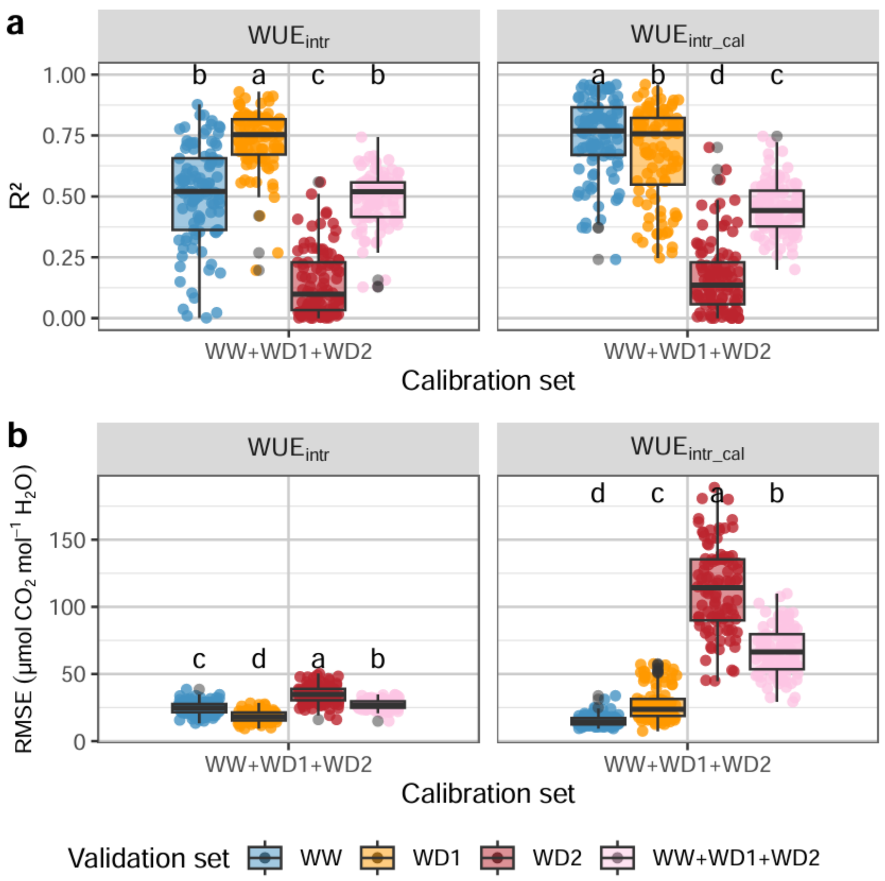


**Figure S7: Prediction quality of PLSR models for $WUE_{intr}$ and $WUE_{intr\_cal}$ depending on the water treatment validation set in the greenhouse environment.**

Models were calibrated with all plants from the three water treatment (WW+WD1+WD2 calibration set) with variables from the Li600 device for intrinsic water use efficiency ($WUE_{intr}$ as the ratio of Li6800 $A_n$ and $g_s$ measurements) directly predicted or calculated $WUE_{intr\_cal}$ as the ratio between the prediction of $A_n$ and the Li600 measurements of $g_s$. **a** corresponds to the $R^2$, **b** corresponds to the RMSE expressed in µmol $CO_2$ $mol^{-1}$ $H_2O$. Each point in the boxplot corresponded to one of the 100 repetitions of the prediction. The blue dots and boxes correspond to the validation set of well-watered treatment (WW); orange dots and boxes to the validation set of moderate water deficit treatment (WD1); and red dots and boxes to the validation set of severe water deficit treatment (WD2).
